# A trade-off shaping leaf mesophyll architecture emerges from a stochastic geometry model

**DOI:** 10.64898/2026.11.18.767598

**Authors:** Milad Rahimi-Majd, Aleca M. Borsuk, Hosein Masoomy, Nancy C. Walker, Craig R. Brodersen, Zoran Nikoloski

**Affiliations:** Bioinformatics Department, Institute of Biochemistry and Biology, University of Potsdam, 14476 Potsdam, Germany; Systems Biology and Mathematical Modeling Group, Max Planck Institute of Molecular Plant Physiology, 14476 Potsdam, Germany; New York Botanical Garden, 2900 Southern Boulevard, 10458 Bronx New York, US; School of the Environment, Yale University, New Haven, CT, 06511 USA

## Abstract

Leaf mesophyll architecture has a great impact on plant functions, including photosynthesis. Yet, understanding the principles governing growth, stability, and interspecific variation of leaf mesophyll architecture remains elusive. Here, we develop a computational framework based on stochastic space tessellation to simulate two-dimensional paradermal slices of leaf spongy mesophyll across 40 vascular plant species using three data-derived parameters: leaf porosity, cell arm length, and cell arm diameter. The framework successfully reconstructs leaf slices while preserving six biological, topological, and cell-level traits, with one- to three-armed cell geometries generally outperforming circular cells. Importantly, we further develop a theoretical model based on stochastic geometry that closely agrees with the simulations and reveals that leaf porosity strongly influences cell wall construction cost, while cell-arm statistics shape this relationship. Our parsimoniously parameterized model provides a basis for optimizing leaf architecture to maximize carbon dioxide uptake while minimizing cell wall construction costs.

## 1 Introduction

Plant leaves constitute the primary site of photosynthesis. Within the leaf, the mesophyll is the main photosynthetic tissue that is generally differentiated into two regions: the palisade mesophyll, composed predominantly of densely packed, cylindrical cells, and the spongy mesophyll, characterized by less densely packed cells that range from spherical to lobed [1]. In addition to cells, the mesophyll contains an extensive network of intercellular air space (IAS) between the cells. This airspace facilitates gas exchange between the atmosphere and mesophyll cells [2]. As a result, the IAS contributes to shaping the leaf mesophyll conductance (*g*_*m*_) that quantifies the ease with which carbon dioxide (*CO*_2_) moves from stomata to its site of fixation by Rubsico in the chloroplast stroma [3]. Further, IAS contributes to modulating both the gas phase component of conductance (*g*_*IAS*_) directly, via diffusional pathways through the IAS, as well as the liquid phase component of mesophyll conductance (*g*_*LIQ*_) indirectly, via diffusional pathways through the mesophyll cell surface area exposed to the IAS [4, 5]. Since mesophyll conductance is regarded as one of the major limitations for photosynthesis [6], understanding the extent to which IAS architecture determines mesophyll conductance can be used as an entry point to optimizing photosynthesis. Specifically, altering the leaf IAS density and the pattern of its distribution has been empirically shown to improve photosynthetic efficiency in plants, unveiling the direct importance of leaf IAS and cell packing geometry to leaf photosynthesis [7]. Beyond its role in carbon assimilation, IAS architecture substantially affects water-use efficiency and plant resilience to changing climatic conditions, making it an important target for crop improvement aimed at stability of food production systems [8].

Despite this essential role of IAS in leaf function, growth, and energy production, the mechanisms governing its development, evolution, and structural stability remain poorly understood [8]. Recent studies have proposed hypotheses regarding the formation and organization of the leaf mesophyll IAS from both dynamic and static perspectives: (i) the developmental mechanisms underlying the emergence and evolution of IAS during cell wall formation, cell division, and epidermal growth [8, 9, 10, 11]; and (ii) the structural and topological principles governing the spatial organization of IAS in mature leaves across plant species [1, 12, 13, 14].

Theories of IAS development in plants posit that it is formed following the establishment of a continuous cellular tissue and arise through three principal developmental mechanisms [8]: (i) lysigeny, involving the generation of tissue cavities via programmed cell death; (ii) schizogeny, in which IAS are created through the physical de-attachment of neighboring cells; and (iii) expansigeny, whereby existing air spaces are expanded through localized expansion of cell wall regions bordering the intercellular space [8, 9]. While lysigeny is considered to be uncommon in plant leaves, the other two processes are considered as the main mechanisms governing leaf IAS expansion, initiating from multicellular junctions [8, 9, 15]. Live-cell imaging of the microtubule cytoskeleton in developing mesophyll cells has provided experimental support for this view, revealing a limited contribution of schizogeny and a predominant role for expansigeny in airspace expansion [16]. More specifically, during an initial schizogeny, the cell wall is divided into free walls (exposed to IAS) and shared walls (cell-cell contact walls), and then almost all subsequent expansion occurs based on the selective expansion of the free walls through expansigeny, while the shared walls remain fixed in size [11].

Alongside the development of computational frameworks for simulating plant cell morphogenesis [17, 18, 19], cell growth based on expansigeny has been modeled using deformable polygons governed by physical laws [10]. The incorporation of key mechanical parameters, *i*.*e*., cell wall stiffness, cell–cell adhesion, and constant boundary pressure, has enabled systematic investigation of the mechanical balance required for spongy mesophyll tissue to remain stable as intercellular void space increases [10].

Following mesophyll development through expansigeny, the final extent of free walls, corresponding to the mesophyll cell surface area exposed to the IAS (*S*_*m*_), plays a critical role in regulating the *CO*_2_ diffusion pathway during *g*_*m*_. This exposed surface forms the interface at which *CO*_2_ transitions from the gaseous phase within the IAS to the liquid phase inside mesophyll cells, potentially affecting both conductance phases *g*_*IAS*_ and *g*_*LIQ*_ [4, 5, 6, 20, 21]. *As a result, an increase in S*_*m*_ enlarges the available surface for *CO*_2_ uptake that can enhance the internal availability of *CO*_2_ within the mesophyll [2, 7, 20]. In contrast, although shared walls are essential for maintaining the mechanical stability of mesophyll tissue [10], an increase of their size incurs additional carbon and resource costs for cell wall construction and maintenance while providing no direct benefit to the gas-exchange interface [22].

In parallel with the IAS dynamic growth models, quantitative analyses of mesophyll tissue architecture have provided new insights into the structural properties of the spongy mesophyll, *i*.*e*., the lower mesophyll layer beneath the palisade mesophyll, characterized by spherical to lobed cells. This layer is of particular interest due to its structural variability and its close association with stomatal pores, which are predominantly located on the lower leaf surface [1, 8, 10, 11]. In particular, recent work has shown that the leaf spongy mesophyll exhibits species-specific structural order, clarifying traditional descriptions of the tissue as largely irregular and non-ordered (1 and references therein). While leaf spongy mesophyll varied along a spectrum of phenotypes, two cell size–shape parameters, *i*.*e*., cell arm length and cell arm diameter, successfully classified 39 vascular plant species into two primary structural classes: honeycomblike and non-honeycomblike. The honeycomb-like species exhibit highly ordered spongy mesophyll organization with prismatic, hexagonally-dominant IAS voids, whereas the latter category corresponds to a less ordered, isotropic cellular network [1]. Importantly, the honeycomb-like structures were shown to follow Euler’s law, revealing a fundamental topological constraint underlying the organization of spongy mesophyll.

Beyond broad geometric and topological classifications of the spongy mesophyll, its underlying geometric principles remain largely unexplored. These principles are key to understanding the constraints governing the expression of leaf morphological traits. Such an understanding could reveal the extent to which individual traits can vary between plant species while maintaining structural stability of the mesophyll tissue.

Here, building on previous findings, we develop the SCellPack framework based on random space tessellation to reconstruct the 2D structure of the paradermal slices of the spongy mesophyll using only three parameters: mesophyll porosity, cell arm length, and cell arm diameter. SCell-Pack successfully reproduces the structural properties of the original slices across several traits. We formulate our model within the framework of a homogeneous Boolean model [23], providing a mechanistic link between free cell wall organization, leaf porosity, and cell arm size. This frame-work identifies the balance between leaf porosity and mean cell arm size, as a key determinant of tissue structure and stability. Our findings demonstrate that a simple stochastic geometry model based on a Poisson point process [23] can capture the geometric organization of the spongy mesophyll across diverse plant species and cell geometries. Finally, using our framework, we derive a data-based relationship between cell wall construction cost and leaf porosity; we then investigate alternative cell morphologies that could shift this relationship, highlighting possible strategies for improving the leaf architecture to maximize *CO*_2_ uptake while minimizing cell wall construction costs.

## 2 Results

### 2.1 SCellPack provides a simulation framework for leaf 2D structures

To model the geometry of the leaf mesophyll, we developed the stochastic cell-packing, SCellPack, framework based on a stochastic space-tessellation approach. We considered five distinct cell geometries: circular, 1-armed, 2-armed, 3-armed, and 4-armed (Fig. 1), with an additional 2-armed geometry having an arm angle of 180^*°*^ (2-armed 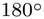) considered in selected analyses (see section 4.2). This definition was motivated by previous work that identified spherical and triply armed morphologies as representative geometries of plant spongy mesophyll cells across 39 plant species [1]. Specifically, the cells were best characterized either by circular (in 2D) geometry or by lobed geometries based on the tissue skeleton with an average of three arms [1]. Building upon this model, we extended the range of considered morphologies to include cells with both fewer and more arms, thereby enabling the investigation and comparison of additional geometrical configurations (see Ref. [1]).

**Figure 1:**
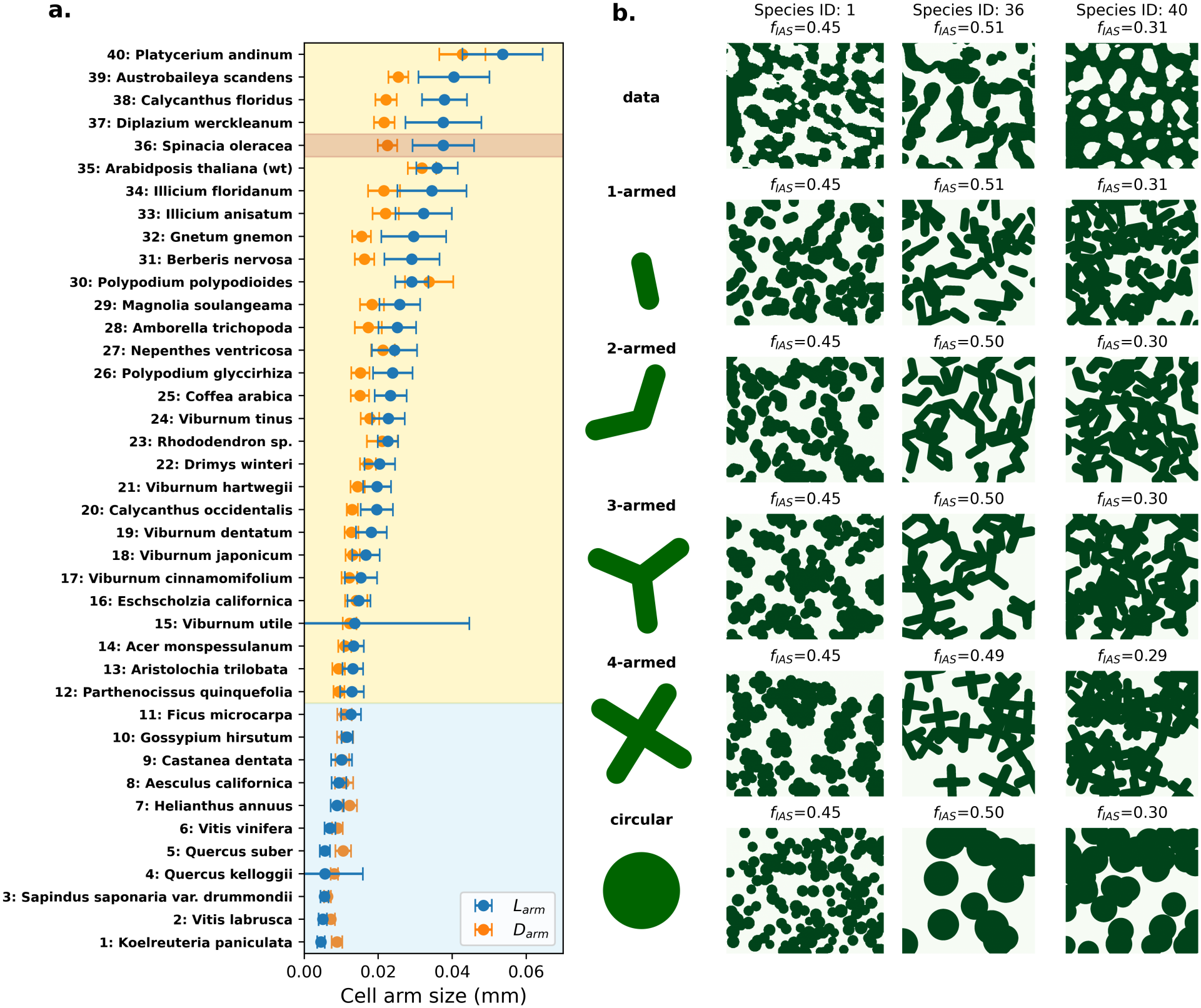
Comparative illustration of simulation results from the SCellPack framework with real-world 2D spongy mesophyll patterns. a. Mean cell arm length (*L*_*arm*_) and mean arm diameter (*D*_*arm*_) of the spongy mesophyll across 40 plant species, with error bars indicating the corresponding standard deviations (see Ref. [1]). Species are ordered by increasing arm length, with blue, yellow, and brown shaded regions corresponding to non-honeycomb, honeycomb, and neither spongy patterns, respectively. **b**. The SCellPack reconstruction for a representative slice of one species from each species category. The selected non-honeycomb and honeycomb species correspond to those with the most extreme mean arm lengths, with the smallest and largest *L*_*arm*_, respectively. The first row shows binary heatmaps of 2D paradermal leaf slices from the three representative species, with cell regions shown in green and intercellular airspace (IAS) in white. The fraction of intercellular airspace to total area (*f*_*IAS*_) and the mesophyll cell surface area exposed to the IAS per unit area (*S*_*m*_) are reported in the panel titles. The lower rows show stochastic simulations generated using SCellPack based on five cell geometries (circular, 1-armed, 2-armed, 3-armed, and 4-armed configurations).

We used two-dimensional paradermal slices from X-ray micro-computed tomography (microCT) images of 40 vascular plant species as the basis for our reconstruction framework (see section 4.1). The simulation box for each species matched the dimensions of the corresponding microCT slice. The fraction of intercellular airspace to total area (porosity; *f*_*IAS*_) of each paradermal slice, together with the leaf-averaged cell arm length (*L*_*arm*_) and arm diameter (*D*_*arm*_), served as input parameters for the simulations. For each cell, values of *L*_*arm*_ and *D*_*arm*_ were randomly sampled from their respective empirical distributions, characterized by the mean and standard deviation of the measured values. Using these parameters, cells were sequentially positioned at random within the simulation domain, with each cell assigned an independently sampled size. Whenever a newly generated candidate cell’s random placement may result in overlaps with one or more previously placed cells, the cell region that could not be located in free area was clipped such that the shared cell walls followed the boundaries of the existing cells, while the free cell wall remained unconstrained. The simulation stops when the target *f*_*IAS*_ (see Fig. 1) or a stopping criterion is reached (see below). This procedure effectively captures the geometrical consequences of expansigeny, but does not provide a dynamic growth model.

Following the classification of the previous study [1], the 40 species were grouped into three categories: non-honeycomb, honeycomb, and neither honeycomb nor spongy (Fig. 1a). The non-honeycomb (11 species) and honeycomb (28 species) categories are characterized by circular (in 2D) and armed cell geometries, respectively (Fig. 1a). The latter category contains only a single species, characterized by elongated cell arms radiating in different directions [1].

We first assessed the integrity of the SCellPack reconstructions by determining whether the framework accurately reproduced the prescribed target parameter. Specifically, because the slice porosity (*f*_*IAS*_) was defined as a target variable based on experimental measurements, its accurate reproduction was a prerequisite for the validity of subsequent analyses (Fig. S1b). The deviations from the target *f*_*IAS*_ may arise from the stopping criterion of the cell-packing procedure, which terminates the simulation after a predefined maximum number of consecutive unsuccessful placement attempts (*i*.*e*., 200), thereby limiting computational cost (section 4.2). We therefore evaluated reconstruction performance across all defined cell geometries, initially considering a sample slice from the representative species within each of the three species categories. No substantial deviations were observed between the porosity of the data paradermal slices and that of the corresponding SCellPack-reconstructed slices. This is supported by the low leaf-level average absolute deviation (≤ 0.03) between the porosity of the data paradermal slices and that of the corresponding SCellPack-simulated images relative to the range of leaf-level average *f*_*IAS*_ values across species (0.24–0.67) (Fig. S1). The negligible deviation can be explained by the substantial free space remaining even at the minimum *f*_*IAS*_: approximately 25% of the simulation domain remained unoccupied in the densest cases. Because the different cell geometries exhibited comparable cell areas, as they are generated from the same cell arm input parameters, this available space was sufficient to reach the prescribed *f*_*IAS*_ across geometries.

The degree of visual agreement between the reconstructed and original microCT images varied across species categories and cell geometries. Consistent with the classification of Borsuk et al. (2022) [1], circular cells showed high visual similarity for non-honeycomb species but substantial differences for species characterized by armed cells (Fig. 1). For instance, 4-armed cells showed weak visual similarity across the representative species, whereas 1-, 2-, and 3-armed cell geometries produced reconstructions that more closely resembled the original microCT images, particularly for non-honeycomb species (Fig. 1). We note that these representative slices are shown only for qualitative illustration, while quantitative evaluation across all species and cell geometries is provided in the following sections.

### 2.2 SCellPack reconstructs the 2D leaf mesophyll cell area exposed to the intercellular airspace

Mesophyll cell surface area exposed to the IAS (*S*_*m*_), which forms the interface between the intercellular airspace and mesophyll cells, is an essential determinant of leaf structure, linking the three-dimensional organization of mesophyll cells to the efficiency of *CO*_2_ diffusion and uptake. [2, 7, 20]. Given the critical importance of *S*_*m*_, this trait was our primary focus for assessing the performance of SCellPack. In the standard leaf physiology measurements, the leaf surface area exposed to the airspace measures the 3D surface area of the cells normalized by the leaf area (*i*.*e*., see Ref. [6]). Since our data consist of 2D paradermal leaf segments, we measured the total exposed cell perimeter at the IAS interface and normalized it by leaf area in the same manner as for the conventional *S*_*m*_ measurements. We retained the conventional notation *S*_*m*_ for this slice-level parameter, which represents the mesophyll cell perimeter exposed to the IAS per unit leaf area.

We first evaluated the extent to which the SCellPack-generated reconstructions reproduced the observed slice-to-slice variation in *S*_*m*_ across individual paradermal slices from three representative species, one from each of the three categories (i.e., honeycomb, non-honeycomb, and neither), using each of the five cell geometries (Fig. 2). Considering the total variation range across the three species, our results showed strong agreement between the measured and reconstructed *S*_*m*_ for the 1-armed, 2-armed, 3-armed, and circular cell geometries across individual slices of the non-honeycomb species with the smallest arm length (*i*.*e*., *Koelreuteria paniculata*), with maximum deviations of 13.0%, 24.1%, 29.3%, and 20.8%, respectively, relative to the total measured *S*_*m*_ variation range across the three species (Fig. 2a). The 4-armed geometry exhibited the greatest deviation, with a maximum value of 38.9%, although the reconstructed values remained consistent with the measured variation (Fig. 2a). In contrast, the two species (*i*.*e*., *Spinacia oleracea* and *Platycerium andinum*) representing the other categories showed strong agreement across all slices for the armed cell geometries (1-, 2-, 3-, and 4-armed), with maximum deviations of 8.9% and 3.7%, respectively, relative to the total *S*_*m*_ variation range across the three species. However, in both cases, reconstructions based on the circular geometries displayed a small but consistent deviation across all slices, with maximum values of 18.9% and 8.7%, respectively (Fig. 2b,c).

**Figure 2:**
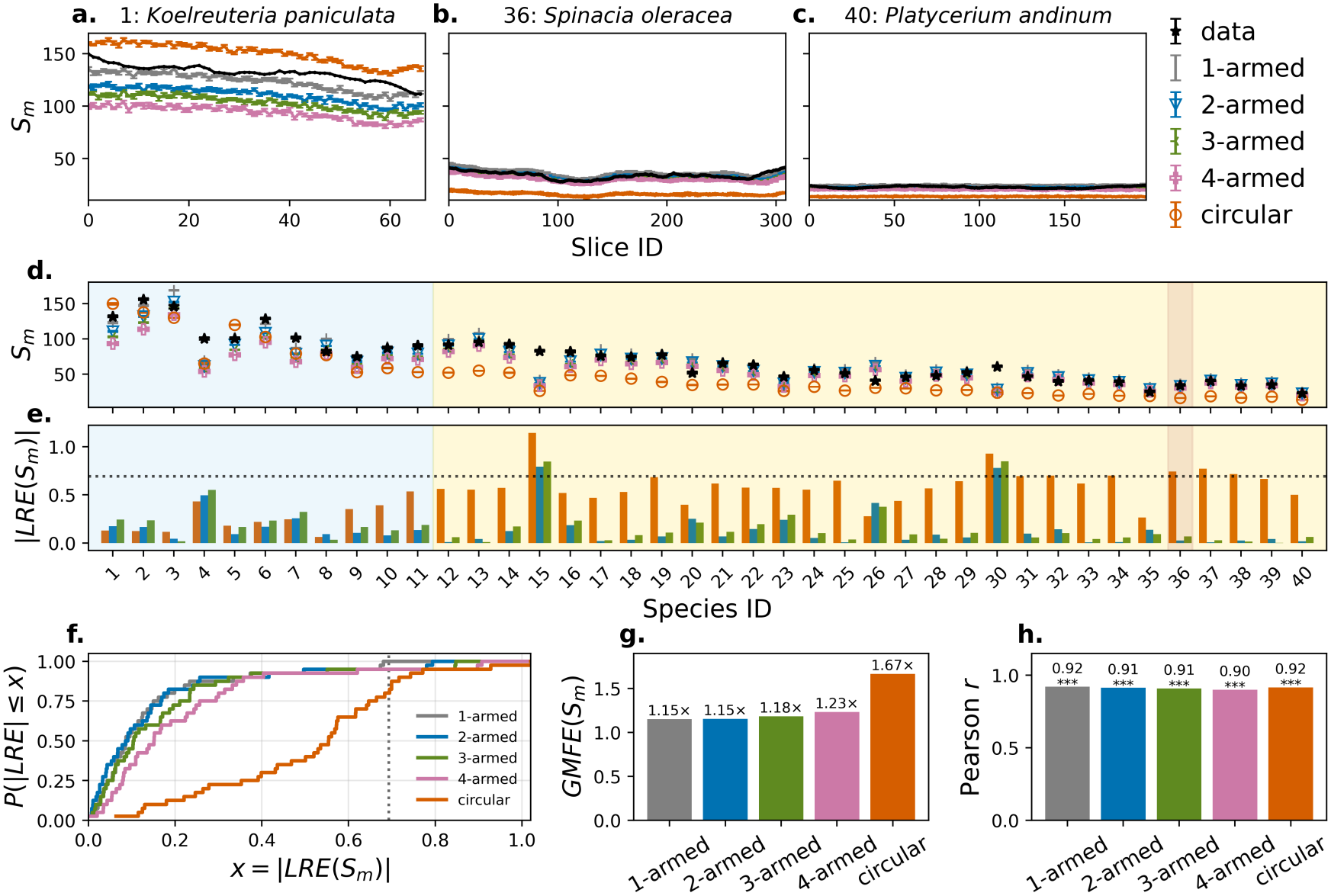
Performance of SCellPack for reconstruction of mesophyll cell perimeter exposed to intercellular airspace (*S*_*m*_). a–c. Slice-level variation in *S*_*m*_ for three representative species, comparing experimental data (black) with SCellPack simulations using five cell geometries (i.e., circular, 1-armed, 2-armed, 3-armed, and 4-armed), marked with different colors (see legend). Error bars indicate the standard errors across the ten independent replicates of the simulations. **d**. Comparison of measured and simulated mean *S*_*m*_ across 40 plant species for all five cell geometries. Error bars indicate the standard errors of the slice-level values. Species are ordered by increasing mean arm length. Blue, yellow, and brown shaded regions indicate non-honeycomb, honeycomb, and neither spongy patterns, respectively. **e**. Absolute logarithmic relative error of *S*_*m*_ (|*LRE*(*S*_*m*_) |), across the 40 species for circular, 2-armed, and 3-armed geometries. Bars indicate the reconstruction errors for the corresponding data points of each geometry in panel **d**. The horizontal dotted line marks a two-fold deviation, |*LRE*| = ln(2). **f**. Cumulative distribution functions of |*LRE*(*S*_*m*_) | for all five cell geometries, showing the probability that the absolute log-relative error is below a given threshold *x*. The vertical dotted line marks a two-fold deviation, |*LRE*| = ln(2). **g**. Geometric mean fold error, *GMFE*(*S*_*m*_), for each cell geometry across the 40 species. **h**. Pearson correlation coefficient, *r*, between simulated and observed mean *S*_*m*_ values across species for each cell geometry. Asterisks indicate statistical significance of the correlation (\*\*\**p <* 0.001). Markers and colors corresponding to the experimental data and simulated cell geometries are consistent across panels.

Next, we assessed these patterns across all 40 species by averaging the measured and reconstructed *S*_*m*_ values across all slices, yielding a single representative value for each species. We quantified the reconstruction error between observed and average simulated *S*_*m*_ values using the absolute logarithmic relative error (|*LRE*(*S*_*m*_)|), a symmetric and scale-invariant metric that quantifies the fold deviation between simulated and observed values (see section 4.5). The analysis showed strong agreement between the measured and reconstructed *S*_*m*_ values across species for all five cell geometries (Fig. 2d). Importantly, however, the magnitude of the reconstruction error differed among cell geometries. Using a two-fold deviation, |*LRE*(*S*_*m*_)| *>* ln(2), as a threshold for acceptable agreement, no species exceeded this threshold for the 1-armed geometry, whereas two species exceeded it for each of the 2-armed, 3-armed, and 4-armed geometries. These errors were concentrated in the same two species across the three mentioned armed-shape geometries.

By contrast, eight species exceeded the two-fold threshold for the circular geometry. Notably, all eight of these species belonged to categories other than non-honeycomb, whereas the circular geometry performed relatively well for non-honeycomb species (Fig. 2d–f). This observation is consistent with the previous classification of non-honeycomb species based on their approximately circular 2D cell shapes. Furthermore, because *D*_*arm*_ *> L*_*arm*_, the armed cell geometries also closely approximated circular cells in these species, resulting in relatively short arms (see the visual patterns in Fig. 1b), thereby explaining their successful reconstruction performance in non-honeycomb species [1].

To obtain a summarized view of the reconstruction performance of each cell geometry across all species, we analyzed the cumulative distributions of |*LRE*(*S*_*m*_)| for each geometry. All 40 species were reconstructed within a two-fold deviation for the 1-armed geometry, whereas 95% of species fell within this range for each of the 2-armed, 3-armed, and 4-armed geometries. In contrast, only 80% of species were reconstructed within a two-fold deviation using the circular geometry (Fig. 2e–f). These results indicate that the armed cell geometries generally provided more accurate reconstructions than the circular geometry, with the 1-armed geometry showing the highest proportion of species reconstructed within two-fold deviation.

We further calculated the geometric mean fold error (*GMFE*) to summarize the typical multiplicative deviation between reconstructed and measured *S*_*m*_ values across species (see Section 4.5). The 1-armed and 2-armed geometries showed the lowest error, with *GMFE* = 1.15× for both geometries, followed by the 3-armed (1.18×) and 4-armed (1.23×) geometries. By contrast, the circular geometry exhibited a substantially higher error (*GMFE* = 1.67×; Fig. 2g).

Finally, we assessed the extent to which each cell geometry captured the interspecific variation in *S*_*m*_ using the Pearson correlation coefficient for the simulated and observed values across species. We found that the correlations were consistently strong and significant across all five geometries, with Pearson’s *r* ranging from 0.90 to 0.92 (Fig. 2h).

Taken together, the individual-slice comparisons and the across-species analyses indicate that SCellPack can reproduce both the variation and magnitude of *S*_*m*_ across a diverse set of species. While all cell geometries showed strong correlations with the measured values, the reconstruction accuracy depended on cell geometry, with the 1-armed and 2-armed geometries consistently showing the lowest errors and the circular geometry showing the poorest overall performance.

### 2.3 Reconstructions based on SCellPack preserve the leaf topology

Beyond reproducing the values for *S*_*m*_, as a geometric property, we next examined whether the SCellPack-generated image reconstructions also preserved the topology of the spongy mesophyll in paradermal slices. To this end, we quantified the number of connected components (*β*_0_), corresponding to the zeroth Betti number, a topological invariant [24, 25], in both the cell phase 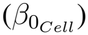 and the IAS phase 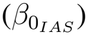 (see section 4.4). These measures capture the connectivity of both phases: the former is relevant to the organization of shared and free cell walls, whereas the latter is an important determinant of *CO*_2_ transport through the IAS in the spongy mesophyll (see Section 1). Because the two phases are spatially complementary, connectivity in one phase also reflects, at least qualitatively, the degree of fragmentation and enclosure in the other, corresponding to more complex structural patterns of the spongy mesophyll.

The SCellPack framework showed good agreement in reconstructing the connectivity of both the cell and IAS phases across the tested cell geometries (Fig. S2). For 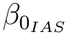 , the non-circular geometries showed particularly strong agreement, with 90–92.5% of species reconstructed within a two-fold deviation and *GMFE* values ranging from 1.36 to 1.38× (Fig. S3a and Table 1). By contrast, the circular geometry showed substantially lower agreement, with only 27.5% of species within a two-fold deviation and a *GMFE* of 2.86×. Consistent with these results, the non-circular geometries showed strong and significant correlations between simulated and observed 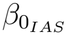 values, with Pearson *r* ranging from 0.69 to 0.72 (*p <* 10^*−*6^). The circular geometry showed a weaker, although still significant, correlation (*r* = 0.55, *p <* 0.001), indicating that circular geometries were moderately effective at reproducing interspecific variation in IAS connectivity. The reduced performance of circular geometries for 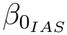 was particularly pronounced among honeycomb-like species. Conversely, circular-cell reconstructions showed relatively good agreement for 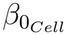 among non-honeycomb species, consistent with their structural organization [1] (Fig. S2).

**Table 1:** SCellPack performance in reconstructing leaf traits. Geometric mean fold error (*GMFE*) and Pearson correlation coefficient (*r*), with *p*-values shown in parentheses, for five leaf traits reconstructed using five cell geometries (circular, 1-armed, 2-armed, 3-armed, and 4-armed) across 40 species.

|  | β<sub>0,*Cell*</sub> |  | β<sub>0,*IAS*</sub> |  | *N*<sub>*Cell*</sub> |  | *P*<sub>*Cell*</sub> |  | *A*<sub>*Cell*</sub> |  |
|---|---|---|---|---|---|---|---|---|---|---|
|  | *GMFE* | r (p) | *GMFE* | r (p) | *GMFE* | r (p) | *GMFE* | r (p) | *GMFE* | r (p) |
| 1-armed | 1.91× | 0.57 (< 0.001) | 1.38× | 0.72 (< 0.001) | 1.61× | 0.63 (< 0.001) | 1.18× | 0.97 (< 0.001) | 1.58× | 0.55 (< 0.001) |
| 2-armed | 1.54× | 0.71 (< 0.001) | 1.38× | 0.71 (< 0.001) | 1.38× | 0.64 (< 0.001) | 1.30× | 0.97 (< 0.001) | 1.41× | 0.52 (< 0.001) |
| 3-armed | 1.74× | 0.76 (< 0.001) | 1.37× | 0.70 (< 0.001) | 1.60× | 0.64 (< 0.001) | 1.56× | 0.97 (< 0.001) | 1.67× | 0.51 (< 0.001) |
| 4-armed | 1.79× | 0.78 (< 0.001) | 1.36× | 0.69 (< 0.001) | 1.86× | 0.64 (< 0.001) | 1.82× | 0.96 (< 0.001) | 1.95× | 0.50 (0.001) |
| Circular | 1.54× | 0.75 (< 0.001) | 2.86× | 0.55 (< 0.001) | 1.89× | 0.58 (< 0.001) | 1.40× | 0.95 (< 0.001) | 1.96× | 0.17 (0.296) |

For 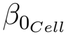, the agreement was more variable across cell geometries, with 60–80% of species reconstructed within a two-fold deviation and *GMFE* values ranging from 1.54 to 1.91× (Fig. S3b and Table 1). The 2-armed geometry showed the highest proportion of species within two-fold (80%) and a low *GMFE* (1.54×), whereas the 1-armed geometry showed the largest *GMFE* (1.91×). Nevertheless, all five geometries captured significant interspecific variation in 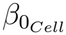, with Pearson *r* ranging from 0.57 to 0.78 (*p <* 0.001). The strongest correlations were observed for the 4-armed (*r* = 0.78), 3-armed (*r* = 0.76), and circular (*r* = 0.75) geometries, whereas the 1-armed geometry showed the weakest correlation (*r* = 0.57).

Overall, these results indicate that the SCellPack framework robustly reproduces connectivity patterns in the leaf spongy mesophyll and captures their interspecific variation, particularly when using armed cell geometries.

### 2.4 SCellPack reconstructions preserve cell statistics based on watershed segmentation

We next evaluated the ability of SCellPack to reconstruct single-cell statistics. Specifically, we assessed the extent to which the reconstructed images reproduced the mean cell number per slice (*N*_*Cell*_), mean cell area per leaf area (*A*_*Cell*_), and mean cell perimeter (*P*_*Cell*_) per leaf area, averaged across all paradermal slices for each species. Cell segmentation was performed using the watershed algorithm applied to the binary images (see section 4.6). The accuracy of the watershed segmentation was evaluated by comparing it with manual segmentation of a representative grayscale paradermal slice for each species (Fig. S6). Considering the variation range across the manually segmented species, watershed segmentation showed high agreement with manual segmentation at the leaf-averaged level, particularly for *A*_Cell_ and *P*_Cell_ (Fig. S7). Therefore, as all subsequent analyses were performed using leaf-averaged statistics, watershed (rather than manual) segmentation was used to quantify the single-cell parameters throughout this study.

The reconstruction of single-cell statistics showed greater variation in performance across cell geometries than observed for *S*_*m*_ and the topological descriptors (Fig. S4). For *N*_*Cell*_, the 2-armed geometry showed the highest proportion of species reconstructed within a two-fold deviation (87.5%), followed by the 3-armed (85.0%) and 1-armed (80.0%) geometries (Fig. S5). The corresponding *GMFE* values were lowest for the 2-armed geometry (1.38×), followed by the 3-armed (1.60×), 1-armed (1.61×), 4-armed (1.86×), and circular (1.89×) geometries (Table 1). Pearson correlations between simulated and observed cell number were significant for all geometries (*r* = 0.58–0.64, *p <* 0.001), with the strongest correlations for the 2-, 3-, and 4-armed geometries (*r* = 0.64) and the weakest for the circular geometry (*r* = 0.58) (Table 1).

For *P*_*Cell*_, the 1-armed geometry showed the strongest agreement, with all species reconstructed within a two-fold deviation (100%), followed by the 2-armed (95.0%) and 3-armed (90.0%) geometries (Fig. S5). These geometries also showed low *GMFE* values, ranging from 1.18× for the 1-armed to 1.56× for the 3-armed geometry (Table 1). Agreement remained relatively high for the circular geometry (92.5% within two-fold; *GMFE* = 1.40×), but was substantially lower for the 4-armed geometry (62.5% within two-fold; *GMFE* = 1.82×) (Fig. S5 and Table 1). Nevertheless, Pearson correlations were consistently strong across all geometries (*r* = 0.95–0.97, all *p <* 0.001), indicating that all geometries captured interspecific variation in mean cell perimeter well (Table 1).

Moreover, *A*_*Cell*_ showed a similar, although more variable pattern. The 2-armed geometry provided the best agreement (85.0% within two-fold; *GMFE* = 1.41×), followed by the 3-armed (82.5%; *GMFE* = 1.67×) and 1-armed (80.0%; *GMFE* = 1.58×) geometries (Fig. S5 and Table 1). The 4-armed and circular geometries showed substantially worse agreement, with only 55.0% of species within two-fold and *GMFE* values of 1.95× and 1.96×, respectively (Fig. S5 and Table 1). Pearson correlations were significant for all armed geometries (*r* = 0.50–0.55, *p* ≤ 0.001), whereas the circular geometry showed no significant correlation (*r* = 0.17, *p* = 0.296) (Table 1). Thus, unlike the armed geometries, the circular geometry was notably less effective at reconstructing cell area and capturing its interspecific variation. However, its reconstruction performance was relatively better among non-honeycomb species (Fig. S4), consistent with the structural organization discussed above.

Overall, the armed geometries generally provided the most consistent reconstruction of single-cell statistics, with the 2-armed geometry showing particularly strong performance across *N*_*Cell*_ and *A*_*Cell*_, while the 1-armed geometry performed best for mean *P*_*Cell*_.

### 2.5 Stochastic geometric model explains the performance of SCellPack

Since our developed model, SCellPack, is based on a random tessellation process, it can be related, at least in part, to existing models and theoretical frameworks in stochastic geometry. The Boolean model (also known as the Poisson germ–grain model) [23] is a particularly relevant framework in this regard, given its random placement of grains (here, cells) within a defined spatial domain. Specifically, the Boolean model describes a random closed set in which the expected fraction of space not covered by the random set, *ρ*, is given by

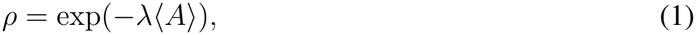

where *λ* is the intensity of the homogeneous Poisson process of germ centers, corresponding to the mean number of germs per unit area, and ⟨*A*⟩ is the mean area of the grains (cells) [23]. In the same line, the specific perimeter of the closed set, *i*.*e*., the boundary length of the union of the grains per unit area of the underlying plane, is given by

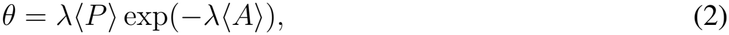

where *λ*⟨*P* ⟩ is the mean perimeter of the grains [26]. Substituting Eq. 1 into Eq. 2, and eliminating *λ*, gives

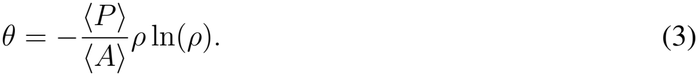

A major difference from our model is that cell placement is sequential in SCellPack. Nevertheless, the porosity and specific perimeter of the resulting cell union are independent of the order in which the cells are placed. As a result, the intercellular air-space fraction (*f*_*IAS*_) and the exposed cell perimeter per unit area (*S*_*m*_) can be regarded as statistical counterparts of the Boolean model quantities *ρ* and *θ*, respectively. This correspondence motivated us to derive an analytical solution for *S*_*m*_ in the SCellPack model based on equation 3, using the data-based mean cell perimeter, mean cell area, and air-space fraction (*i*.*e*., porosity) as inputs.

However, deriving an analytical expression for *S*_*m*_ in SCellPack directly from the Boolean model is not straightforward because of differences in the implementation of cell placement. In SCellPack, candidate cells that are completely overlapped by previously placed cells are rejected (see Section 4.2), which may introduce some deviations from the assumptions of the Boolean model. The major challenge, however, arises from the clipping of cells when they partially overlap with previously placed cells (see Section 4.2). Clipping changes the area and perimeter of individual cells relative to their initially sampled values and consequently modifies the resulting cell-area and cell-perimeter distributions. Importantly, however, clipping does not affect the porosity or specific perimeter of the resulting structure. This is because clipping occurs within the union of the cells and therefore does not alter either the area or perimeter of the outer boundary of the union. Therefore, the model can still be characterized using the mean cell area and perimeter prior to clipping.

Accordingly, here we rely on the arm statistics rather than on the final, truncated cell sizes. The initial cell sizes, prior to clipping, are directly determined by the distributions of the arm statistics. For each cell geometry, the mean perimeter and area of the initial, non-truncated cells can therefore be obtained analytically from the moments of the arm-length and arm-diameter distributions. Specifically, one can derive ⟨*A*⟩ and ⟨*P* ⟩ from the distributions of *L*_*arm*_ and *D*_*arm*_, while accounting for the error terms associated with the sampling of these parameters. In this line, to assess the performance of the Boolean model, we derived the corresponding expressions for two fundamental geometries, circular and 1-armed cells, as well as for the 2-armed 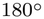 geometry. The selection of 2-armed 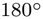 was motivated by its representation as a simple extension of the 1-armed cell, while consistently showing excellent agreement with the primary 2-armed geometry (characterized by an arm angle of 120^*°*^) in reconstructing the previously assessed parameters (see Fig. S2 and Fig. S4). These analytical derivations are presented in Section 4.7. Analytical solutions for more complex cell geometries, involving additional arms and different arm angles, require further extensions of these relations. A detailed derivation for such geometries is left for future work.

The formulation of the Boolean model, based on the initial cell sizes rather than the final truncated sizes, also ensures that the Boolean model uses exactly the same inputs as SCellPack, *i*.*e*., 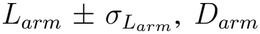, and *f*_*IAS*_. We refer to the values of *ρ* and *θ* calculated from these analytical solutions as the theoretical *f*_*IAS*_ and *S*_*m*_, respectively (see above).

Importantly, *S*_*m*_ calculated from the Boolean model showed excellent agreement with the corresponding values obtained from the SCellPack simulations across all three cell-shape geometries (Fig. S8 and Fig. 3). For all 40 species (100%), the theoretical values of *S*_*m*_ were within two-fold of the SCellPack results for each geometry. The agreement was further supported by highly significant Pearson correlations for the circular (*r* = 0.998), 1-armed (*r* = 0.997), and 2-armed 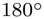 (*r* = 0.997) geometries. Consistent with these strong correlations, the *GMFE* error values were very low, at 1.03×, 1.04×, and 1.06× for the 1-armed, 2-armed, and 2-armed 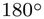 geometries, respectively (Fig. 3). Thus, the Boolean model predictions closely reproduced the SCellPack-derived *S*_*m*_ values across all three geometries, and the deviations between theory and data closely followed the same patterns as the deviations between the SCellPack-derived *S*_*m*_ values and the empirical data described above (Section 2.2; Fig. S8 and Fig. 3).

**Figure 3:**
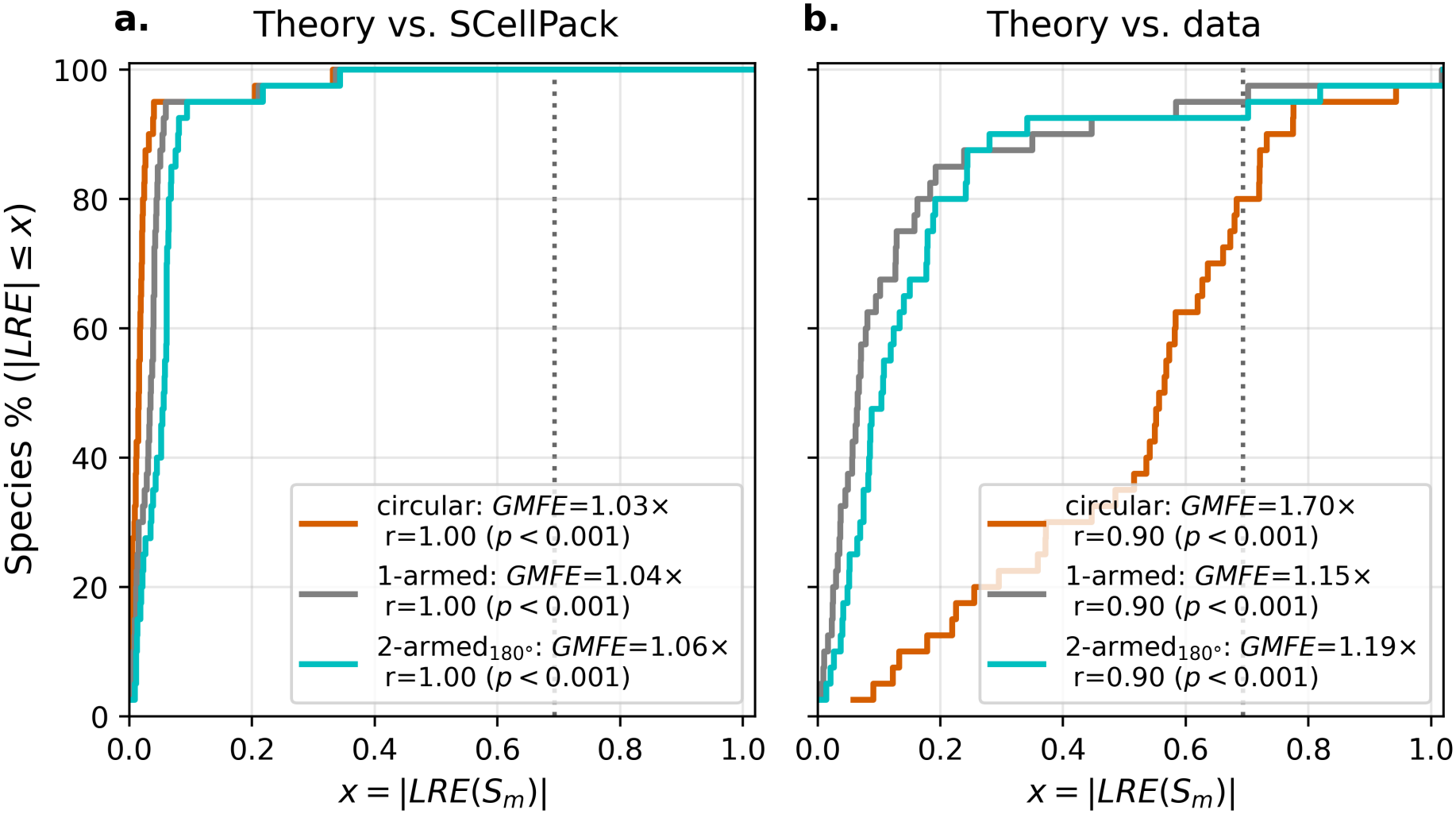
Performance of the Boolean model in estimating mesophyll cell surface area exposed to intercellular airspace (*S*_*m*_). Cumulative distribution functions of the absolute logarithmic relative error |*LRE*| between: **a**. theoretical Boolean model and SCellPack simulated values of *S*_*m*_, and **b**. theoretical Boolean model and data measured values of *S*_*m*_. Colors indicate different cell geometries (circular, 1-armed, 2-armed 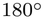). The vertical dashed lines mark |*LRE*| = ln(2), corresponding to 2× fold-error threshold.

Together, this formulation analytically links cell arm statistics and leaf porosity to cell wall properties, providing a simple stochastic-geometric framework for describing how these structural features shape leaf architecture as captured by SCellPack.

### 2.6 The economical geometry for optimal cell-airspace gas-exchange

Beyond the importance of *S*_*m*_ for leaf *CO*_2_ conductance, the balance between maximizing free cell wall, as captured by *S*_*m*_, and minimizing shared cell walls is another aspect that plays an essential role in determining the structural costs associated with cell wall construction (see section 1). To quantify this cost trade-off, we define the loss fraction as

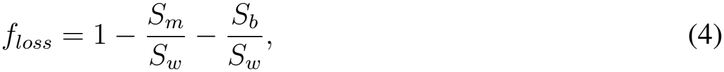

where *S*_*b*_ and *S*_*w*_ denote the cell perimeter exposed to the image border and the total cell wall perimeter, respectively, both normalized by leaf area, analogous to *S*_*m*_. We note that *S*_*b*_ introduces a deviation from the real biological system due to the artificial image boundaries, and its relative contribution diminishes as the image size increases. The defined quantity, *f*_*loss*_, represents the fraction of the cell wall that does not contribute to IAS gas exchange because it is located at cell–cell interfaces, and therefore constitutes an additional structural cost of cell wall construction.

Based on the Boolean model, using *S*_*b*_ = 0, *S*_*w*_ = *λ*⟨*P* ⟩, along with equations 1, 2, and 3, while treating *ρ*, and *θ* as the counterparts of *f*_*IAS*_, and *S*_*m*_, respectively, we obtain

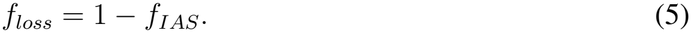

This relation indicates a perfect linear trade-off between leaf porosity and the fraction of shared cell walls predicted by the Boolean model (see Section 2.5).

Whilst the Boolean model treats overlapping regions between cells as valid cell area, SCell-Pack prevents cell overlap, consistent with biological constraints. By definition, this results in shorter shared wall lengths due to cell truncation, *i*.*e*., a smaller *S*_*w*_, while maintaining consistent *S*_*m*_. Interestingly, we observed an excellent correlation between *f*_*loss*_ and (*f*_*IAS*_) across the SCell-Pack reconstructions generated using the different cell geometries (Fig. 4a-e). Specifically, the SCellPack reconstructions exhibited very strong relationships, with coefficients of determination consistently high across all cell geometries (*R*^2^ ≥ 0.96) (Fig. 4a-e). The fitted relationships were also remarkably consistent across geometries, with slopes of −0.88, −0.88, −0.87, −0.86, and −0.87 for the 1-, 2-, 3-, 4-armed, and circular geometries, respectively (Fig. 4a-e). These findings suggest that, despite differences in cell geometry, the SCellPack reconstructions are in line with the linear trade-off between *f*_*loss*_ and *f*_*IAS*_, predicted by the Boolean model.

**Figure 4:**
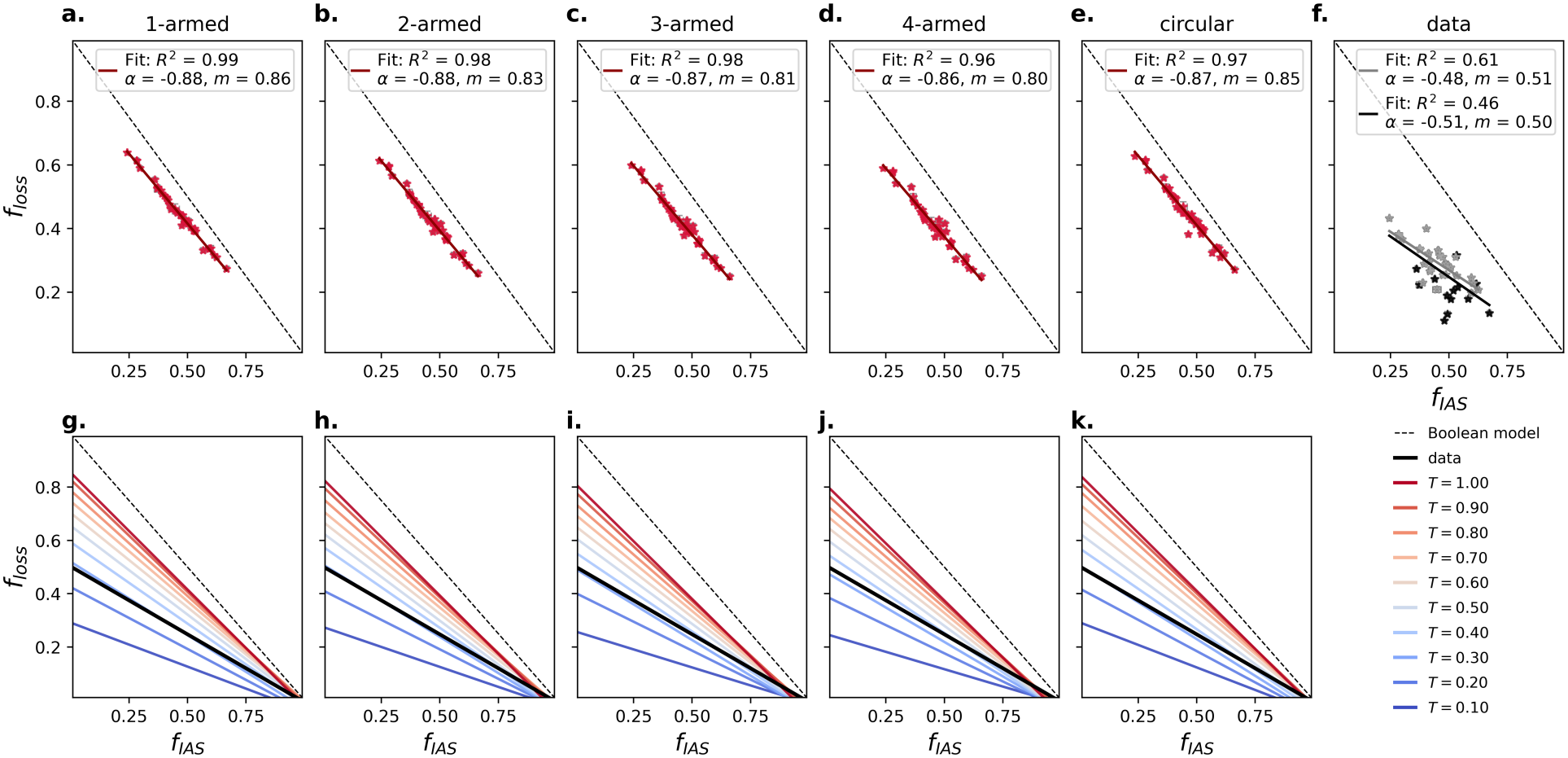
Association between leaf porosity and cell shared wall fraction. Linear fits of the shared-wall fraction (*f*_*loss*_) as a function of the intercellular airspace fraction (*f*_*IAS*_) for **a**. 1-armed, **b**. 2-armed, **c**. 3-armed, **d**. 4-armed, and **e**. circular cell geometries, and **f**. measured data. The red data points and red lines in panels **a-e** represent the SCellPack results at the truncation fraction of *T* = 1, while the black data points and black fit in panel **f** represent the measured data; the gray data points and fit in panel **f** show the subset of honeycomb species. The lower panels **g-k** correspond to panels **a-e**, respectively, and are aligned in the same columns, showing the corresponding linear fits across the range of *T* tested in SCellPack. In panels **g-k**, colored lines represent the linear fit for each truncation fraction, the black line represents the fit to the measured data, and the dashed line indicates the ideal trade-off predicted by the Boolean model. The vertical and horizontal error bars in panels **a-f** represent the standard errors of the mean values across paradermal slices for *f*_*loss*_ and *f*_*IAS*_, respectively. In the upper panels, *R*^2^, *α*, and *m* denote the coefficient of determination, slope, and intercept of the linear fits, respectively.

Similarly, we observed a moderate to strong relationship between *f*_*loss*_ and *f*_*IAS*_ in the measured data across all as well as only for the honeycomb species. Specifically, the measured data across all 40 species showed a coefficient of determination of *R*^2^ = 0.46, with a substantially shallower slope of −0.51 and an intercept of 0.50 (Fig. 4f). Restricting the analysis to honeycomb species strengthened the relationship, yielded *R*^2^ = 0.61, with a similar slope of −0.48. These results indicate a more pronounced deviation from the ideal linear trade-off observed in the Boolean model for the measured data, while highlighting *f*_*IAS*_ as an important determinant of the costs associated with cell wall construction.

The higher slopes and intercepts observed for the SCellPack reconstructions indicate that this framework remains closer to the assumptions of the Boolean model than the measured data in terms of the total cell perimeter. This is the case since both predict a substantially larger total cell perimeter relative to *S*_*m*_ (Eq. 4). One possible explanation for this observation is that SCellPack considers even very small non-overlapping portions of candidate cells as valid cells, which can generate additional small cells and increase the total cell perimeter (see Section 4.2). To test this hypothesis, we introduced an additional constraint through the clipping fraction parameter (*T*), which defines the upper-bound fraction of a candidate cell that can be truncated by previously placed cells (see Section 4.8).

We tested and compared *T* values in the range [0.1, 1.0] with a step of 0.1, where *T* = 1.0 corresponds to the default SCellPack setting, with no truncation constraint except rejection of candidate cells with complete overlap (sections 4.2 and 4.8). The SCellPack-generated results showed strong linear relationships across all tested *T* values, with *R*^2^ ≥ 0.87 at *T* = 0.1 and *R*^2^ ≥ 0.95 for *T >* 0.1 (Fig. S9). Importantly, both the slope and intercept of the linear *f*_*loss*_-*f*_*IAS*_ fits decreased systematically with decreasing *T* , following similar trends across cell geometries (Fig. 4g-k and Fig. S9). Moreover, depending on cell geometry, the SCellPack-fitted relationships most closely matched the measured data around *T* = 0.3 − 0.4 (Fig. 4g-k and Fig. S9).

To assess the ability of SCellPack to reconstruct leaf structural parameters using the *T* value that provided the best agreement with the experimental *f*_*loss*_–*f*_*IAS*_ relationship, we next evaluated all previously assessed traits, namely *S*_*m*_, *β*_0,*IAS*_, *β*_0,*Cell*_, *N*_*Cell*_, *P*_*Cell*_, and *A*_*Cell*_, using SCellPack configured with *T* = 0.4. Among the cell geometries, the 2-armed cells consistently showed comparable or even higher reconstruction accuracy across species for all traits relative to the primary cases with *T* = 1, whereas the other geometries showed notable declines in the reconstruction for one or more traits (see Fig. S10). These results support the relevance of incorporating the *T* constraint into SCellPack, particularly for specific cell geometries and structural traits.

In summary, we showed that *f*_*loss*_ is negatively correlated with *f*_*IAS*_ across plant species. This indicates that plant species exhibiting low porosity incur higher cell wall construction costs relative to the free cell wall perimeter available for *CO*_2_ transport. We consistently observed this relationship across the Boolean model, the SCellPack framework, and the measured data, although the measured data exhibited a notably less steep decline in *f*_*loss*_ with increasing *f*_*IAS*_ than predicted by either model. We further demonstrated that varying the truncation fraction *T* changes both the slope and intercept of the resulting linear fits.

Specifically, we observe closely matching patterns in the dependence of the linear fits on *T* across all cell-shape geometries (Fig. 4g-k, S9). Varying the value of *T* modifies the leaf-averaged cell perimeter and area distributions relative to those of the random closed set represented by the Boolean model, while producing similar responses across different cell-shape geometries. Therefore, these results suggest that cell size and area distributions are stronger determinants of cell-construction costs than cell-shape geometry. Together, these findings provide a basis for identifying potential economical strategies to modulate crop improvement by optimizing the balance between leaf *f*_*IAS*_ and cell-size distributions, thereby maximizing *CO*_2_ uptake while minimizing the costs of cell wall construction.

## 3 Discussion

Recent advances in leaf imaging techniques have improved our understanding of the internal structure of the leaf mesophyll. However, the general mechanisms governing the development and maintenance of mesophyll structure remain poorly understood. In particular, the spongy mesophyll exhibits remarkable diversity in the spatial organization of cells and intercellular airspaces (IAS) across vascular plant species, yet the mechanisms underlying the formation and structural stability of these distinct architectures remain largely unresolved [1, 8].

At the same time, improving *CO*_2_ uptake through mesophyll conductance while minimizing the resource costs associated with cell wall construction necessitates a mechanistic understanding of how mesophyll architecture influences these functions. Motivated by this challenge, we developed SCellPack framework, based on stochastic space tessellation, for simulating the two-dimensional architecture of the leaf spongy mesophyll across species. Specifically, SCellPack aims to reconstruct the binary structure, comprising either cell or intercellular airspace phases, of two-dimensional paradermal leaf sections derived from three-dimensional microCT volumes using a small number of data-derived parameters. Building on previous work that categorized the spectrum of spongy mesophyll phenotype into three primary classes based on cell-arm length and diameter [1], we parameterized SCellPack using a single slice-level parameter, leaf porosity (*f*_*IAS*_), and two leaf-level parameters: cell-arm length (*L*_*arm*_) and cell-arm diameter (*D*_*arm*_). This parsimonious parameterization provides a key advantage in terms of model simplicity compared with similar simulation frameworks, which generally involve a larger set of parameters (*e*.*g*., [10, 18, 19]).

Motivated by the study of Borsuk et al. (2022) [1], we configured five cell geometries within the SCellPack framework. In that study, species were categorized according to circular (in 2D) and triply armed cell-shape geometries, which were associated with non-honeycomb and honeycomb-like architectures, respectively, as well as a third category comprising cells with radially elongated arms that conform to neither geometry. Building on this classification, we considered the circular geometry and extended the triply armed cell geometry to include both fewer and greater numbers of arms, considering 1-, 2-, 3-, and 4-armed cell geometries.

The simulation procedure in SCellPack involves truncating candidate cells such that they occupy only the available space on their free-wall sides, while the shared walls formed with previously placed cells remain fixed (see section 4.2). This procedure partially captures, at a coarsegrained level, expansigeny, a widely accepted mechanism proposed to underlie IAS formation [8, 11]. In particular, the truncation criterion imposes a directional constraint on cell expansion that is effectively consistent with the proposed role of expansigeny in shaping the IAS. Thus, although the framework represents the static architecture of mature mesophyll, the resulting cell configurations may effectively provide insights into the mechanical constraints associated with IAS formation and organization.

To assess the performance of SCellPack, we focused primarily on the cell surface perimeter exposed to the IAS in 2D (S_m_), given its critical role in mesophyll conductance (*g*_*m*_) as the interface for *CO*_2_ transfer from the IAS into the cell, as well as its implications for the costs associated with cell wall construction (see section 1). Our results showed excellent agreement between experimentally measured and SCellPack-reconstructed *S*_*m*_ values at both the slice and leaf levels (Fig. 2 and section 2.2). At the leaf level, the performance of SCellPack was assessed using three complementary metrics: the percentage of species for which the reconstructed *S*_*m*_ was within a two-fold error of the measured value (80% for circular and 95% for armed cell-shape species), the geometric mean fold error (*GMFE* = 1.67× for circular and *GMFE* ≤ 1.23× for armed cells), and the

Pearson correlation between measured and simulated values across species (*r* ≥ 0.9 for all cell geometries) (Fig. 2 and section 2.2). Together, these metrics provided complementary evidence of the high representativeness of SCellPack simulations: the high percentage of species predicted within a two-fold error together with the low *GMFE* indicates biologically meaningful simulations at the individual-species level. Moreover, the high Pearson correlation demonstrates the ability of SCellPack to reproduce interspecific variation, independently of the absolute accuracy of individual species-level realizations resulting from SCellPack. Therefore, while simulations from SCellPack result in different realizations, they are highly representative of the analyzed mesophyll architectures.

We next found that SCellPack consistently achieved good reconstruction accuracy across the majority of cases, with performance varying among cell geometries and traits, for the reconstruction of two topological traits (Section 2.3 and Fig. S2) and three cell-level traits (section 2.4 and Fig. S3). The ability to preserve topological traits ensures that the connectivity of the IAS, which is essential for *CO*_2_ diffusion, and the connectivity of cell objects, which is relevant to the regulation of shared and free cell walls, are maintained in the images generated by SCellPack. In addition, the accurate reconstruction of cell-level statistics, *i*.*e*., mean perimeter, area, and cell number, importantly, indicates the biological relevance of cell truncation during cell tessellation in SCellPack framework, in the leaf average level. Overall, the ability of SCellPack to reconstruct leaf paradermal images while preserving six key biological, topological, and anatomical traits demonstrates its capacity to comprehensively reproduce the spatial organization of leaf paradermal slices across a broad range of species representing all previously identified cell-shape categories.

Among the five tested cell geometries, the 1-, 2-, and 3-armed geometries consistently ranked among the best-performing configurations across all the species (sections 1, 2.3, and 2.4). In contrast, circular cells performed acceptably primarily for the 11 non-honeycomb species, while in the majority of cases they failed to adequately reconstruct the slices for the remaining species (Figs. 2, S2, and S4). These findings are consistent with the hypothesis proposed by Borsuk et al. (2022) [1], in which circular cell geometries in 2D were considered the geometric endpoint of triply armed cells as *D*_*arm*_ *> L*_*arm*_. This property is also incorporated into our framework (see Fig. 1), ensuring that armed cells effectively converge towards a circular geometry under this condition, whereas the converse does not hold.

We next showed that SCellPack can in part be captured by the Boolean model, a theoretical framework from stochastic geometry [23]. Specifically, we derived theoretical estimates of *S*_*m*_ from analytical expressions for the mean cell area and perimeter, parameterized by the empirically measured *L*_*arm*_ and *D*_*arm*_ for each cell geometry (section 2.5). Our theoretical model for *S*_*m*_ is a function of *f*_*IAS*_, *L*_*arm*_ (and its standard deviation, 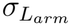), and *D*_*arm*_ (equation 3), the same set of parameters used as inputs to SCellPack. Importantly, we observed excellent agreement between the Boolean model estimates and SCellPack simulations, which in turn showed high agreement with the experimental measurements for three representative cell geometries (Fig. 3), demonstrating the validity of the theoretical formulation and its consistency with the underlying SCellPack framework.

The analytical expressions in our extension of the Boolean model describe the uniform random distributions of candidate cell area and perimeter, parameterized by 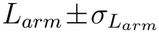 and *D*_*arm*_, equivalent to the cell distributions generated by SCellPack prior to their truncation during the tessellation procedure. Thus, these distributions do not correspond directly to the realized distributions of cell area and perimeter after tessellation, nor to those measured from segmented experimental images. Nevertheless, because *L*_*arm*_ and *D*_*arm*_ are directly measured from experimental data, the high accuracy of the *S*_*m*_ reconstructions obtained with the Boolean model indicates that these pre-tessellation distributions retain biological relevance. Given that cell overlap is permitted in the Boolean model, these distributions may be interpreted as representing the idealized cell size and geometry under unconstrained growth, in the absence of the shared cell wall constraints that arise through expansigeny.

The stability of the spongy mesophyll architecture represents another poorly understood aspect of spongy mesophyll structure [1, 10, 11]. In particular, the structural robustness of the spongy mesophyll despite the presence of a substantial fraction of the mesophyll as void space (*f*_*IAS*_ = 0.24–0.67 in our dataset; see Fig. S1 for details) highlights the need to understand how porosity is regulated by cell structural parameters. Here, our theoretical model provides a mechanistic framework for understanding the regulation of *f*_*IAS*_, indicating that the coordinated regulation of *S*_*m*_, *L*_*arm*_, and *D*_*arm*_ contributes to the maintenance of this architecture, as described by Eq. 3.

Finally, we assessed the proportion of shared cell wall relative to the total cell wall (*f*_*loss*_) as a measure of the structural investment in cell wall construction that is unavailable for *CO*_2_ exchange between the IAS and the cell interior (section 2.6). For the idealized case described by the Boolean model, we showed a perfect linear trade-off between the two fraction parameters, *f*_*loss*_ and *f*_*IAS*_ (*f*_*loss*_ = 1 − *f*_*IAS*_), noting that the model considers the perimeter of overlapping cell regions as shared cell walls (section 2.6). While accounting for the biologically relevant shared walls of truncated cells in SCellPack, simulations nevertheless showed an excellent linear relationship between *f*_*loss*_ and *f*_*IAS*_, with only small deviations in the slope and intercept relative to the ideal trade-off predicted by the Boolean model (Fig. 4a-e). Importantly, the experimental data also showed a moderate (strong) relationship across all (honeycomb-like) species, between *f*_*loss*_ and *f*_*IAS*_, indicating that *f*_*loss*_ is closely associated with leaf porosity partuculrly for the honeycomb-like species (Fig. 4f). Specifically, the results suggest that species with lower porosity exhibit greater cost-effectiveness in cell wall construction compared with species characterized by a higher proportion of IAS.

Despite exhibiting the same *f*_*loss*_–*f*_*IAS*_ correlation trend, the slopes and intercepts of the experimental linear fits were substantially lower than those obtained from the SCellPack framework (Fig. 4). This discrepancy indicates that, in real tissues, the maximum attainable *f*_*loss*_ is lower and that *f*_*loss*_ is less sensitive to variation in *f*_*IAS*_ across species than obtained by the SCellPack simulations. Toward strategies for the optimization of the costs associated with cell wall construction as a function of leaf internal architecture, we explored alternative distributions of truncated cell sizes and shapes in SCellPack. Specifically, we introduced an upper-bound truncation fraction, *T* , ranging from 0 to 1, where lower values of *T* shift the cell population toward its ideal, non-truncated geometry (sections 2.6 and 4.8). This approach modifies the spatial distribution of cells from the uniform Poisson distribution assumed in the Boolean model [23], thereby allowing us to examine how variations in cell size, shape, and spatial organization influence the costs associated with cell wall construction.

We observed that decreasing *T* systematically altered the *f*_*loss*_–*f*_*IAS*_ relationship, with both the slope and intercept gradually decreasing until converging with the experimental data fits at *T* ∼ 0.3–0.4 across all cell geometries (see Fig. 4g-k and Fig. S9). This suggests that the real data sets retain approximately 60–70% of the idealized geometrical characteristics defined by the cell-arm statistics. In addition, the linear fits obtained from the SCellPack simulations across different *T* values exhibited a consistent trend across all five cell geometries (Fig. 4g-k and Fig. S9). This consistency suggests that the distribution of cell-arm statistics plays a dominant role in determining the associated construction costs, whereas variations in mean cell area, perimeter, and overall cell morphology appear to have a comparatively smaller effect across the different cell geometries. Together, these findings suggest that the regulation of leaf porosity with final cell arm statistics determines the degree of cost inefficiency associated with cell wall construction. Moreover, these findings provide potential pathways for cost–benefit optimization of leaf internal architecture across different cell geometries, with the aim of enhancing *CO*_2_ uptake while minimizing the costs associated with cell wall construction.

While SCellPack and the theoretical framework presented here advance our understanding of the geometrical principles underlying leaf tissue architecture and morphogenesis, extending the approach to three-dimensional imaging represents a complementary avenue for future research. The current 2D model could also benefit from incorporating slice-level arm statistics instead of the leaf-level statistics available for and used in the present analysis, potentially improving reconstruction accuracy. Furthermore, both the 2D and 3D models could benefit from inclusion of the spatial distribution of stomatal pores. In particular, sub-stomatal cavities require IAS voids that are connected to the rest of the IAS and have been shown to influence mesophyll and IAS architecture [27, 28]. Extending the formulation to more generalized non-homogeneous Boolean process [23], and other similar sequential space tessellation approaches such as the dead leaves model [29, 30, 31] is another direction that could improve the theoretical explanation of the mesophyll structural patterns. Finally, developing predictive models for porosity and cell-arm statistics from available species data sets could enable their prediction for unseen species and genotypes, which, when integrated into SCellPack, could facilitate the reconstruction of leaf mesophyll architecture across previously uncharacterized species and genotypes.

## 4 Methods

### 4.1 Data set

The data set used in this study consists of X-ray micro-computed tomography (microCT) images of the spongy mesophyll from 40 plant species, originally published by Borsuk *et al*. (2022) [1]. The data set encompasses plant species from 30 genera and 24 families, representing a broad diversity of extant vascular plants [1]. Leaf tissue samples, approximately 1.5-2 mm wide and 15 mm long, were excised near the leaf midpoint, offset 5-10 mm from the mid vein, and imaged at the Advanced Light Source (ALS, Lawrence Berkeley National Laboratory) and the TOMCAT beamline of the Swiss Light Source (SLS). The reconstructed image stacks have pixel resolutions ranging from 0.1625 to 1.277 *µ*m.

We used the two-dimensional paradermal slices extracted from the three-dimensional microCT reconstructions together with the accompanying species-level metadata. These metadata include the pixel resolution and the mean and standard deviation of the cell arm length (*L*_*arm*_) and arm diameter (*D*_*arm*_), which were used to parameterize our analyses.

Each species is represented by a leaf tissue sample with different image dimensions and numbers of available paradermal slices through the spongy mesophyll. To preserve the original structural information, all analyses and simulations were performed using the native image size and spatial resolution of each species. When simulations required varying the analysis domain or tissue depth, these parameters were selected within the intervals defined by the original data set, relative to the dimensions of the imported image stack.

### 4.2 Simulation

Synthetic paradermal binary images were simulated for the complete data set of 40 species, mentioned above. Because species differed in both the dimensions of the paradermal 2D rectangular box and the number of available paradermal slices, simulations were performed independently for each slice using its original image dimensions, ensuring that all geometric measurements remained directly comparable to the original microCT data.

Five idealized cell morphologies were considered: circular cells and cells with one, two, three, or four protruding arms (hereafter referred to as 1-armed, 2-armed, 3-armed, and 4-armed cells, respectively). The characteristic cell size was sampled from the experimentally measured distribution for each species. Specifically, arm length (*A*_*arm*_) and arm diameter (*D*_*arm*_) were independently sampled from normal distributions defined by the measured mean and standard deviation for the corresponding species at the leaf level, as provided in Ref. [1]. For the simulations, these geometric parameters were expressed relative to the simulation box size, thereby preserving the experimentally observed proportions between cell arm dimensions and tissue image dimensions for all species. Each armed cell was constructed as the union of buffered line segments. Specifically, each arm was represented by a line segment of length *L*_*arm*_ originating from a common center and expanded by a circular buffer of radius *D*_*arm*_*/*2, corresponding to the Minkowski sum of the line segment with a circle of radius *D*_*arm*_*/*2. The circular buffer was numerically approximated using 32 segments per quarter circle (Python package *Shapely* version 2.1.2, `buffer` function). Similarly, circular cells were generated by buffering a point at the cell center with radius *L*_*arm*_. This construction produces rounded arm termini and smoothly connected junctions through the union operation, resulting in geometries with well-defined areas and perimeters determined by *L*_*arm*_, *D*_*arm*_, and arm orientations. When *D*_*arm*_ *> L*_*arm*_, the armed morphologies increasingly approached a circular shape. The angles between arms for the 2-, 3-, and 4-armed cell geometries were fixed at 120^*°*^, 120^*°*^, and 90^*°*^, respectively. To assess the effect of arm angle, 2-armed cells with an angle of 180^*°*^ were also simulated and referred to as 2-armed 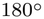 throughout the text. Additionally, each cell was assigned a random orientation during placement.

The simulation was based on a sequential random space-tessellation process in which cells were generated and packed within a rectangular simulation domain. The dimensions of the simulation box were defined from the corresponding image slice dimensions. To maintain consistent scaling between cell size and box size across image slices, the experimentally measured cell arm lengths were normalized by the diagonal of the image slice array, expressed in physical units. Specifically, for each species, the mean (*L*_*arm*_) and standard deviation (*σ*_*L*_) of the arm length were converted to dimensionless quantities as

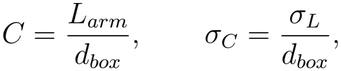

where *d*_*box*_ is the diagonal of the rectangular simulation box. The arm length of each candidate cell was then defined as *c* × *d*_*box*_, where *c* was sampled uniformly from the range *C* ± *σ*_*C*_. To avoid inconsistencies between the sampled cell arm length and arm diameter for an individual cell, the arm diameter was not sampled independently. Instead, for each species, the ratio between the mean arm diameter and mean arm length was calculated as 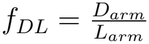. For each sampled cell, the arm diameter was subsequently calculated from its sampled arm length using the scaling parameter *f*_*DL*_. Thus, cells with different sampled arm lengths retained the same ratio of *D*_*arm*_*/L*_*arm*_ within a given species. Each candidate cell, with a randomly sampled size and morphology, was generated at a uniformly random position within the simulation box and tested against previously accepted cells. If the candidate cell overlapped previously accepted cells, the overlapping region was clipped against the boundaries of the existing cells, while the remaining free boundary of the candidate cell was left unconstrained. In some cases, a portion of a candidate cell that did not directly overlap an existing cell could nevertheless become disconnected from the largest remaining portion as a consequence of overlaps with other cells. Such disconnected regions were treated as effectively overlapping areas and removed accordingly. If one or more existing cells completely covered the candidate cell, or if the candidate cell completely covered one or more previously accepted cells, the trial was rejected. Cells intersecting the simulation-box boundary were similarly clipped, using the same area-retention criterion as for cell-cell overlap.

The porosity (*f*_*IAS*_), defined as the fraction of empty area relative to the total image area, was prescribed as the stopping criterion for the cell placement process. Cell placement continued until the accumulated cell area reached the target porosity, at which point the simulation was terminated. Simulations were additionally stopped when a predefined maximum number (*i*.*e*., 200) of consecutive unsuccessful placement attempts was reached, indicating that further cell insertion was not possible under the overlap constraint. This deviation from the target porosity was considered the simulation error, particularly affecting images with low porosity. In all simulations, an ensemble of 10 independent realizations was generated for each species–cell morphology combination. Ensemble statistics were calculated from these realizations to obtain the mean structural properties and their associated variability, represented by the mean standard deviation across realizations.

All structural statistics of the simulated slices were calculated after converting the *Shapely* cell geometries into raster images. Each cell was assigned a unique integer label, with the background assigned zero. The labeled images were analyzed using the same image-based procedure applied to the experimental slices. Specifically, the parameters were initially obtained in raster-pixel units and then converted to the simulation scale using the raster pixel scale. All the parameters with a physical length unit are expressed based on the physical length in milimeters by converting the images based on their pixel size.

### 4.3 Cell surface area exposed to the intercellular airspace

In the standard leaf physiology measurements, the leaf surface area exposed to the airspace is normalized by the leaf area (*i*.*e*., see Ref. [6]). Since here our data are leaf 2D paradermal segments, we measure the total exposed cell perimeter exposed to the airspace (*S*_*m*_) with the same normalization to the leaf area. In these measurements, the cell perimeters were computed using a 4-connected neighborhood with the *scikit-image* Python package (version 0.26.0) [32], using the `measure.perimeter()` function.

### 4.4 Topological parameters

The number of connected components is a topological descriptor of a binary image, corresponding to the zeroth Betti number in topological terms [24, 25]. In two dimensions, it is defined as the number of disconnected objects. Accordingly, two topological descriptors were analyzed: the number of cell connected components (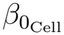) and the number of intercellular airspace (IAS) connected components (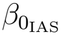). Both phases were labeled using a 4-connected neighborhood, computed with the *SciPy* Python package (version 1.16.3) [33]. This analysis was performed separately for the cell and IAS phases of both the measured and reconstructed paradermal slices.

### 4.5 Reconstruction error metrics

To quantify the deviation between observed, simulated, and theoretically calculated trait values across species, we used the absolute Logarithmic Relative Error (|*LRE*|), defined as

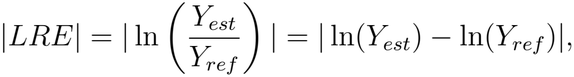

where *Y*_*est*_ and *Y*_*ref*_ denote the estimated and reference trait values, respectively. This metric quantifies the magnitude of the deviation between estimated and reference values on a multiplicative scale, with |*LRE*| = 0 indicating perfect agreement. Throughout the text, we express the deviation as the equivalent fold difference, exp(|*LRE*|), denoted by ×. Specifically, we use a two-fold difference, corresponding to |*LRE*| = 0.693 and exp(0.693) ≈ 2×, as a threshold for assessing reconstruction performance. This threshold indicates that the estimated value is either twice or half the reference value. A key advantage of |*LRE*| for model assessment is its scale invariance. This property is particularly relevant for image-derived parameters, whose values may depend on the choice of spatial units (e.g., pixels versus physical dimensions).

To summarize the |*LRE*| across species, we used the geometric mean fold error (*GMFE*), defined as

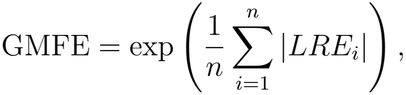

where *i* indexes species and *n* is the number of species. The *GMFE* ranges from 1 to ∞, with values closer to 1 indicating more accurate reconstructions and larger values indicating greater typical fold deviation between estimated and reference values.

### 4.6 Cell segmentation

Cell segmentation was performed using a watershed-based procedure in ImageJ [34] to identify cell boundaries (Fig. S6). Because watershed segmentation removed the cell boundary separating adjacent cells, cell areas and perimeters derived from this procedure included a small associated approximation. To validate the slice-level performance of the watershed segmentation, we manually corrected the segmentation results for a representative grayscale paradermal slice of each species. Specifically, we first manually segmented the intercellular airspace in the grayscale images and then applied the watershed algorithm to the corresponding binarised cell-tissue masks. Next, extraneous cell boundaries were removed or added by comparing the initial watershed output with the grayscale images, and incorrectly assigned void pixels were reassigned to the cell phase, or vice versa, following a similar procedure to that described in Ref. [10]. The accuracy of the watershed segmentation was then evaluated by comparing the mean and standard deviation of the resulting cell statistics with those obtained from the manually corrected segmentations for each species (Fig. S6).

### 4.7 The cell area and perimeter equations based on cell arm statistics

Here we derived the relationships for the expected area and perimeter of the cells prior to any truncation resulting from cell-cell overlap or intersection with the boundaries of the simulation domain in the SCellPack framework.

For the circular geometry, the cell area (*A*_*c*_) and perimeter (*P*_*c*_) of the initial, non-truncated cells, are determined solely by their randomly sampled arm length *L*, considered as the radius (see section 4.2), given by *πL*^2^ and 2*πL*, respectively. Hence, the average values of these parameters over the cells generated by SCellpaCk is given by:

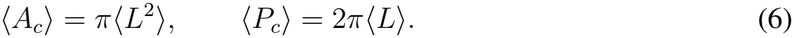

In the simulations, the cell radius *L* is sampled uniformly from the interval [*L*_arm_ − *σ*_*L*_, *L*_arm_ + *σ*_*L*_], where *L*_arm_ and *σ*_*L*_ denote the experimentally measured mean and standard deviation of the arm length, respectively. It is easy to show that the first two moments of this distribution are

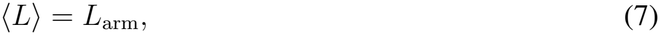

and

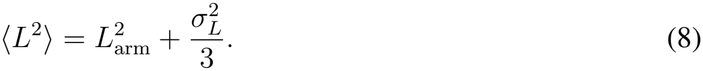

Substituting the equations 7 and 8 into the equation 6, the mean area and perimeter of the initially generated circular cells by SCellPack are

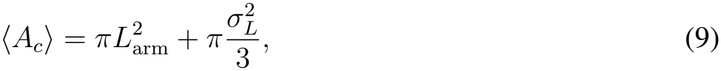

and

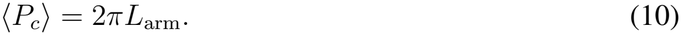

For the 1-armed cell geometry, the cell is constructed as the Minkowski sum (buffer) of a line segment of length *L* by a circular radius *D/*2, where *D* is the corresponding sampled arm diameter. The resulting capsule-like geometry consists of a rectangular section of length *L* and width *D*, with two semicircular ends of radius *D/*2 (see Fig. 1b). Its area and perimeter are therefore given by

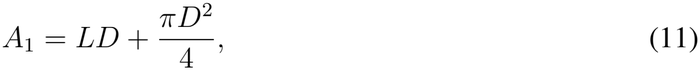

and

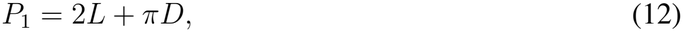

respectively. Here, the first terms in each expression correspond to the rectangular section, while the second terms account for the two semicircular ends. In our simulations, *D* is scaled proportionally to *L* (see section 4.2) as:

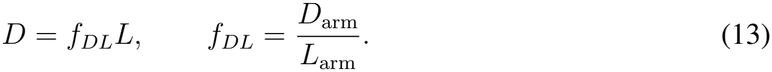

Therefore, substituting the equation 13 into equations 11 and 12 the average area and perimeter over randomly sampled cells are given by:

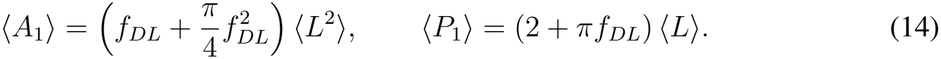

Finally, substituting equations 13, 7 and 8 into equation 14, the mean area and perimeter of the initially generated 1-armed cells by SCellPack become:

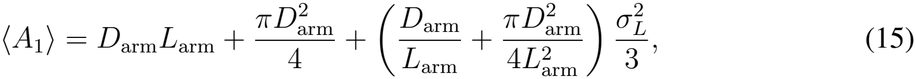

and

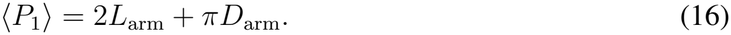

For cells with multiple arms, the area and perimeter depend not only on the arm length and diameter, but also on the angles between the arms. This is because the buffered arms overlap in the region surrounding their common origin. Specifically, for an n-armed cell, the area and perimeter would be the combined armed statistics minus their overlapped sections as

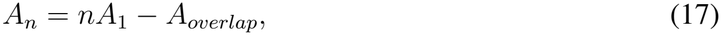

and

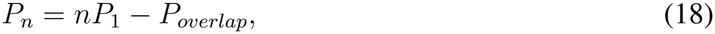

where *A*_*overlap*_ represents the total area correction associated with pairwise and higher-order overlaps, while *P*_*overlap*_ represents the corresponding reduction in the summed individual-arm perimeters due to boundaries that become internal to the union. For instance, for the cell morphology 2-armed 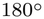 one can easily show that 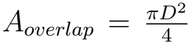 and *P*_*overlap*_ = *πD*, while by substituting these in the previous relations and performing the same procedure, we will have:

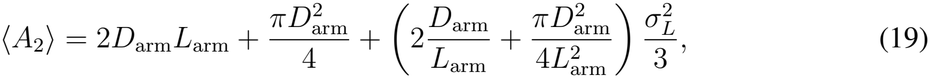

and

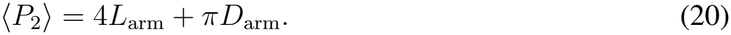

### 4.8 Cell truncation factor

To limit the extent to which the clipping process during the SCellPack simulation can reduce a candidate cell from its initially sampled size, as described above, a cell truncation fraction parameter *T* was introduced. The parameter *T* defines the upper-bound fraction of the originally sampled candidate-cell area that can be truncated due to interactions with previously accepted cells or the simulation-box boundary. Based on this definition, *T* was varied from 0 to 1. A value of *T* = 1 imposes no upper limit on truncation, provided that a non-zero area remains for the candidate cell, whereas *T* = 0 does not allow any truncation of the initially sampled cell; otherwise, the trial is rejected. Smaller values of *T* ensure that the placed cells more closely retain their initially sampled size and prescribed morphology (e.g., circular or 3-armed geometries), but also make it more difficult to reach the target *f*_*IAS*_, possibly resulting in a larger deviation from the target and, consequently, a higher simulation error.

## 5 Code and data availability

The Python functions supporting our analyses are available at the following URL: github.com/MRahimiMajd/SCel All the binary MicroCT slices of the species analyzed in the study are publicly available through the Ref. [1].

## 8 Author contributions

M.R.-M. conceptualized the study, performed computations, analyzed data, interpreted results, and wrote the paper. A.M.B. curated data, interpreted results, and edited the paper. H.M. performed manual segmentation of the grayscale images and contributed software and investigation. N.C.W. contributed data curation and interpretation. C.R.B. interpreted results and edited the paper. Z.N. conceptualized the study, interpreted results, acquired funding, supervised M.R.-M. and H.M., and wrote the paper. All authors contributed to finalizing the manuscript.

## Supporting Information

**Figure S1:**
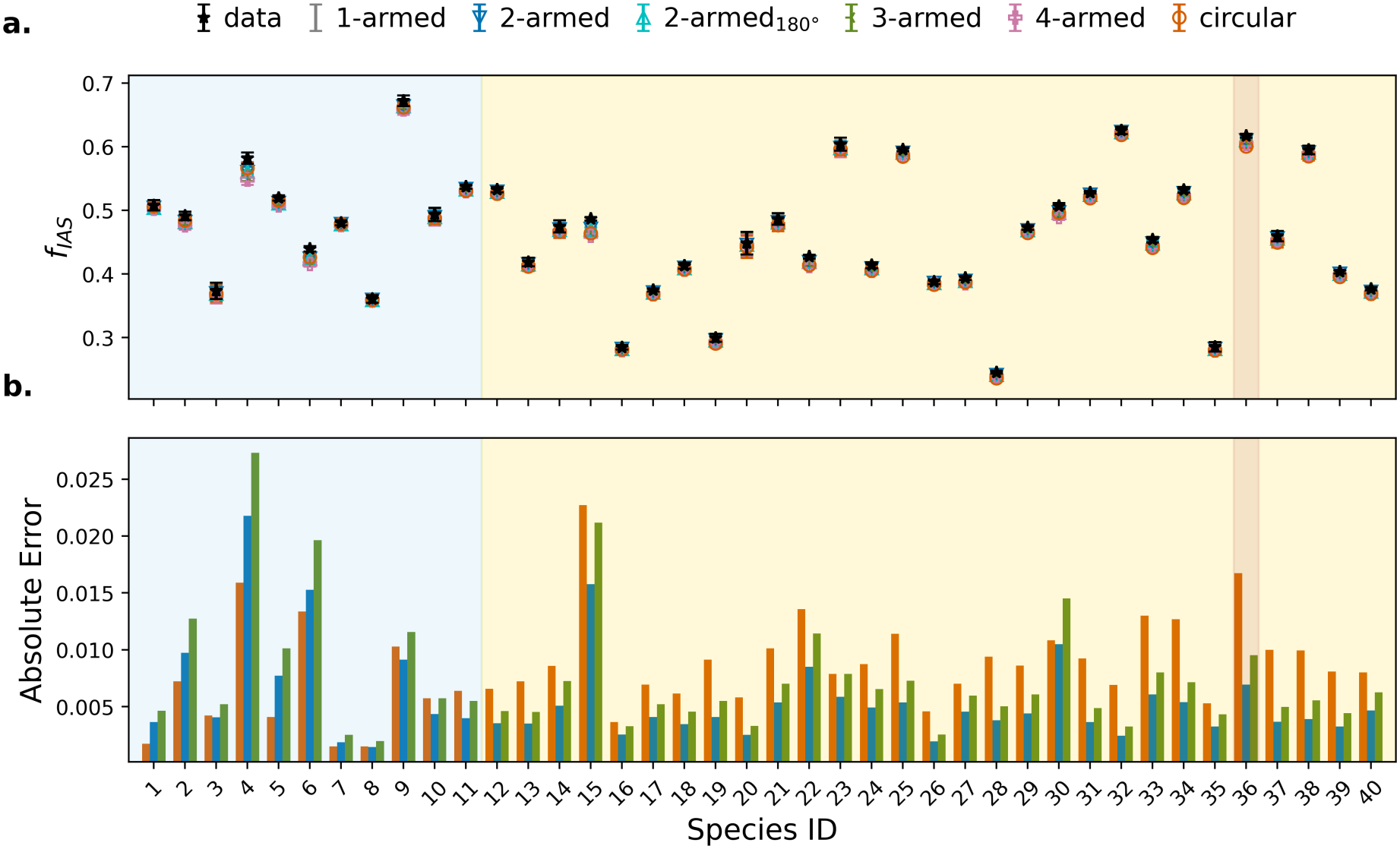
SCellPack statistics for the reconstructed leaf porosity. a. Comparison of the measured and simulated mean fraction of intercellular airspace to total area (*f*_*IAS*_) in paradermal leaf slices across 40 plant species. Each marker corresponds to a different cell geometry (circular, 1-armed, 2-armed, 2-armed 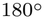, 3-armed, and 4-armed). Error bars indicate standard errors of slice-averaged values. **b**. The bar plot of the absolute error between the measured and simulated values of 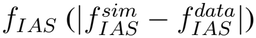 for three selected cell geometries (circular, 2-armed, and 3-armed) across all species. In both panels, species are ordered by increasing arm length, with blue, yellow, and brown shaded regions indicating non-honeycomb, honeycomb, and neither spongy patterns, respectively. Markers and colors are consistent across all panels.

**Figure S2:**
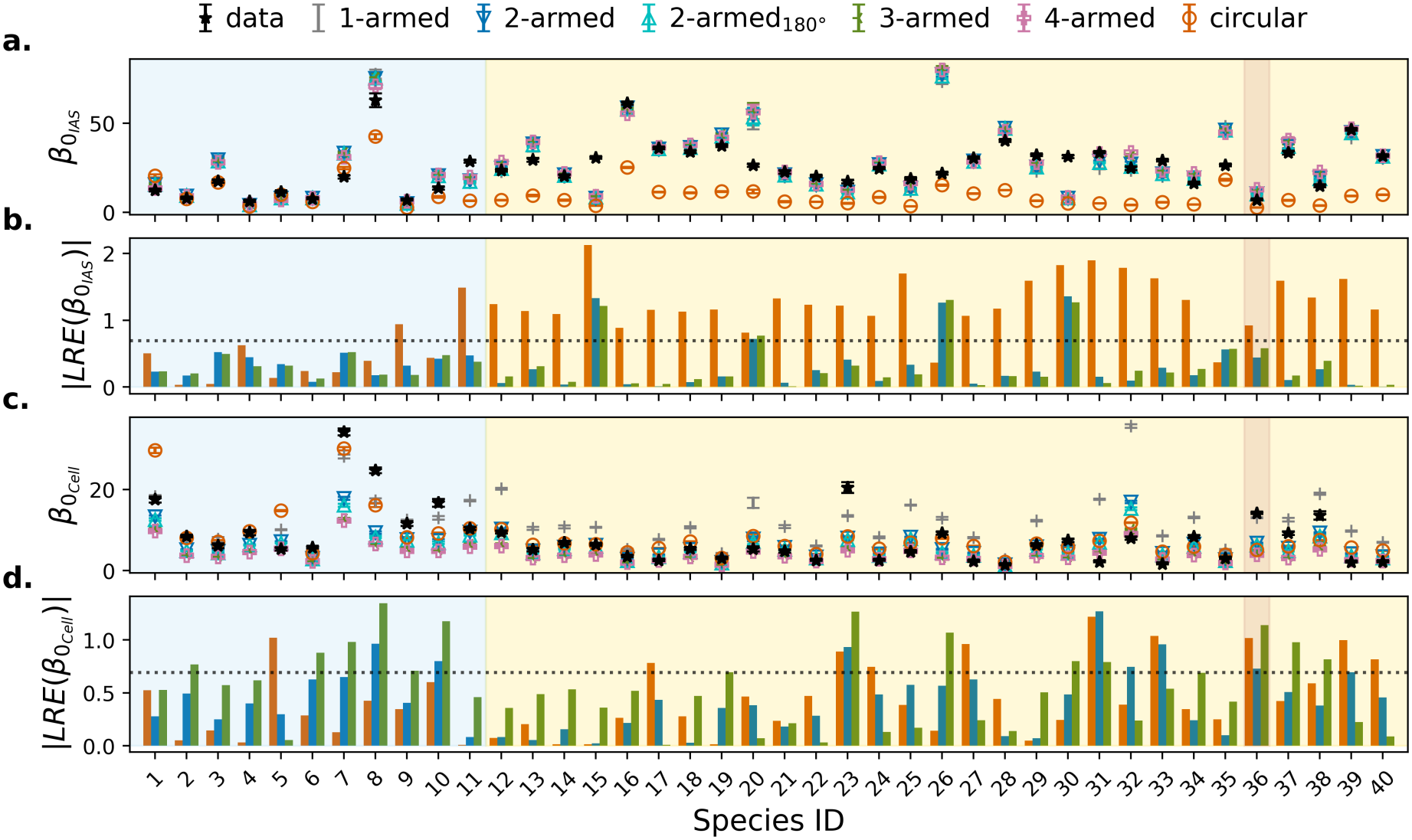
SCellPack statistics for the reconstructed leaf topology traits. a. Comparison between measured and simulated mean number of connected components of leaf intracellular airspace (IAS) phase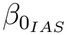 across 40 plant species. Each marker corresponds to a different cell geometry (circular, 1-armed, 2-armed, 2-armed 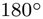, 3-armed, and 4-armed). Error bars indicate the standard errors of the slice-level values. Species are ordered by increasing mean arm length. Blue, yellow, and brown shaded regions indicate non-honeycomb, honeycomb, and neither spongy patterns, respectively. **b**. Absolute logarithmic relative error of 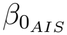 (|*LRE*(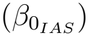) |)) across the 40 species for circular, 2-armed, and 3-armed geometries. Bars indicate the reconstruction errors for the corresponding data points of each geometry in panel **a**. The horizontal dotted line marks a two-fold deviation, |*LRE*| = ln(2). **c**. and **d**. Are the same as **a** and **b** for the number of connected components of leaf cell phase (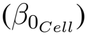), respectively. Markers and colors corresponding to the experimental data and simulated cell geometries are consistent across panels.

**Figure S3:**
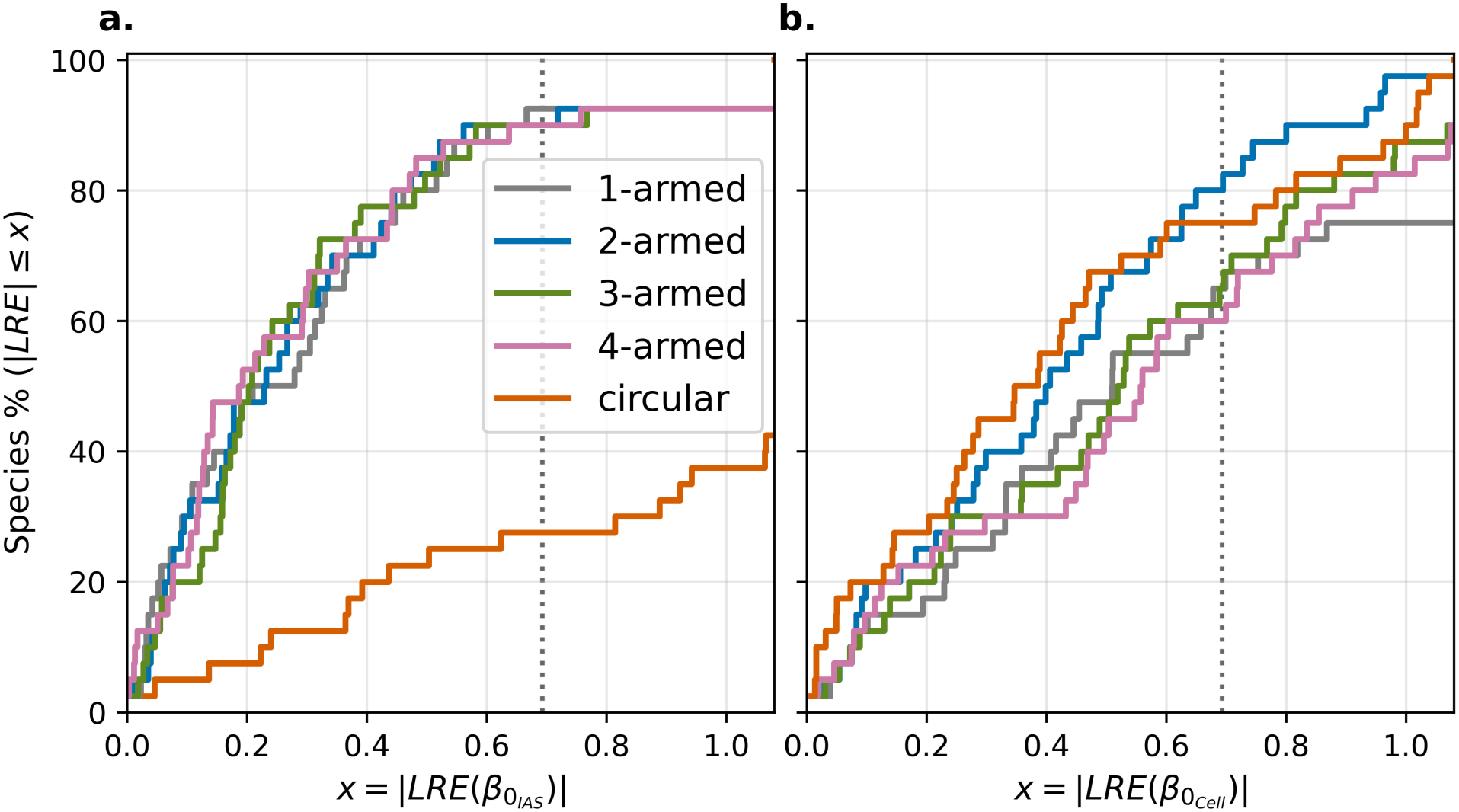
Distributions of SCellPack reconstruction errors for leaf topology. Cumulative distribution functions of the absolute logarithmic error (|*LRE*|) between simulated and measured values for the traits: **a**. number of intercellular airspace (IAS) connected components (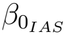) and **b**. number of cell connected components (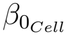). Colors indicate different cell geometries (circular, 1-armed, 2-armed, 3-armed, and 4-armed). The curves show the probability that the reconstruction error is less than or equal to a given threshold *x*. The vertical dotted lines mark a two-fold deviation, |*LRE*| = ln(2).

**Figure S4:**
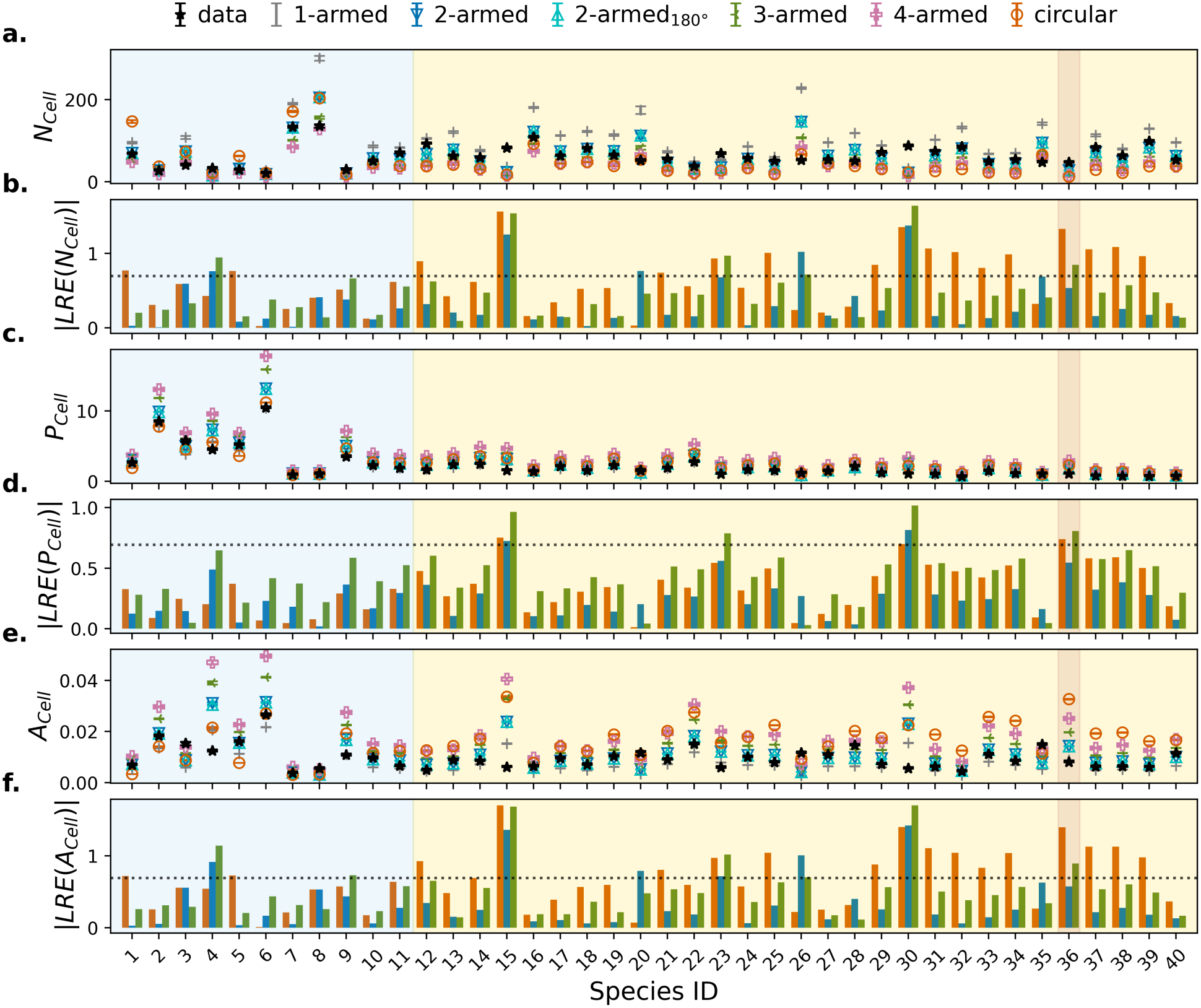
SCellPack statistics for the reconstructed leaf cell-level traits. a. Comparison between measured and simulated mean cell number (*N*_*Cell*_) across 40 plant species. Each marker corresponds to a different cell geometry (circular, 1-armed, 2-armed, 2-armed 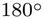, 3-armed, and 4-armed). Error bars indicate the standard errors of the slice-level values. Species are ordered by increasing mean arm length. Blue, yellow, and brown shaded regions indicate non-honeycomb, honeycomb, and neither spongy patterns, respectively. **b**. Absolute logarithmic relative error of *N*_*Cell*_ (|*LRE*(*N*_*Cell*_) |)) across the 40 species for circular, 2-armed, and 3-armed geometries. Bars indicate the reconstruction errors for the corresponding data points of each geometry in panel **a**. The horizontal dotted line marks a two-fold deviation, |*LRE*| = ln(2). **c**. and **e**., are the same as **a** for the average cell perimeter per leaf area (*P*_*Cell*_), and the average cell area per leaf area (*A*_*Cell*_), respectively. **d**. and **f**. are the same as **b** for *P*_*Cell*_ (|*LRE*(*P*_*Cell*_) |) and *A*_*Cell*_ (|*LRE*(*A*_*Cell*_) |), respectively. The measured values in all panels are based on the watershed cell segmentation of leaf paradermal slices. Markers and colors are consistent across all panels.

**Figure S5:**
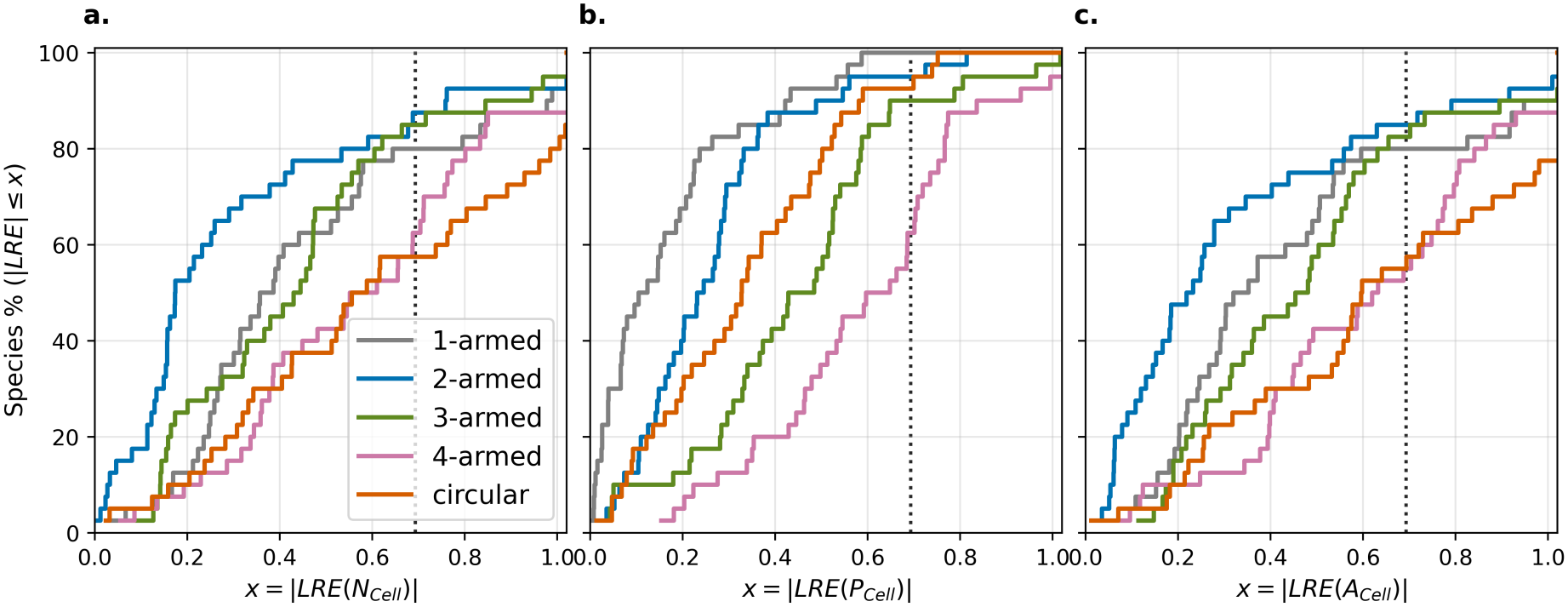
Distributions of SCellPack reconstruction errors for leaf cell-level traits. Cumulative distribution functions of the absolute logarithmic error (|*LRE*|) between simulated and measured values for the traits: a. the number of cells (*N*_*Cell*_), b. the average cell perimeter per leaf area (*P*_*Cell*_), and c. the average cell area per leaf area (*A*_*Cell*_). Colors indicate different cell geometries (circular, 1-armed, 2-armed, 3-armed, and 4-armed). The curves show the probability that the reconstruction error is less than or equal to a given threshold *x*. The vertical dotted lines mark a two-fold deviation, |*LRE*| = ln(2).

**Figure S6:**
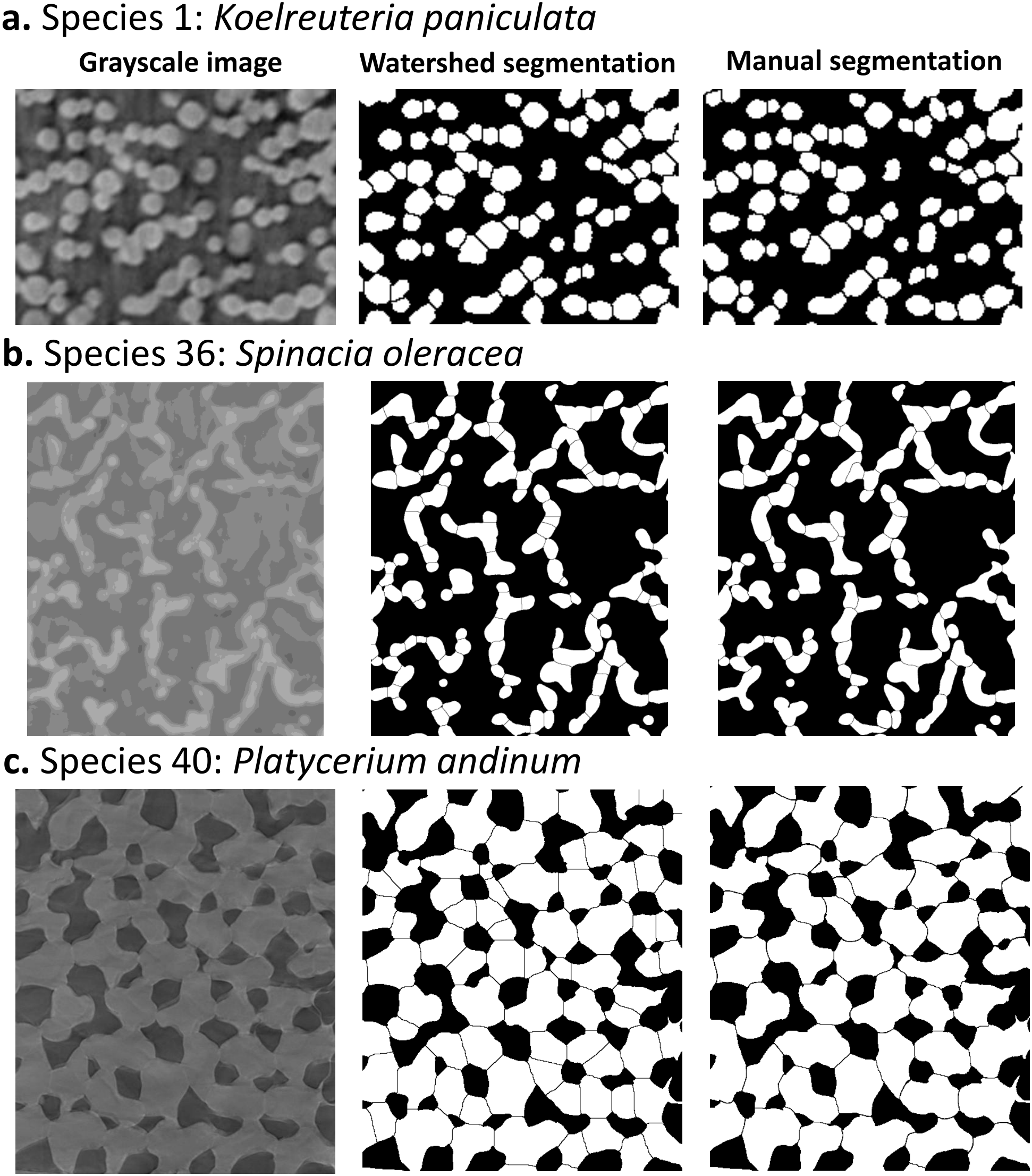
Comparison of watershed and manual cell segmentation of leaf paradermal slices. Segmentation workflow for spongy mesophyll cells from original grayscale microCT images for three representative species. For each species, the left, middle, and right panels show the original grayscale image, the watershed segmentation, and the manually corrected watershed segmentation, respectively.

**Figure S7:**
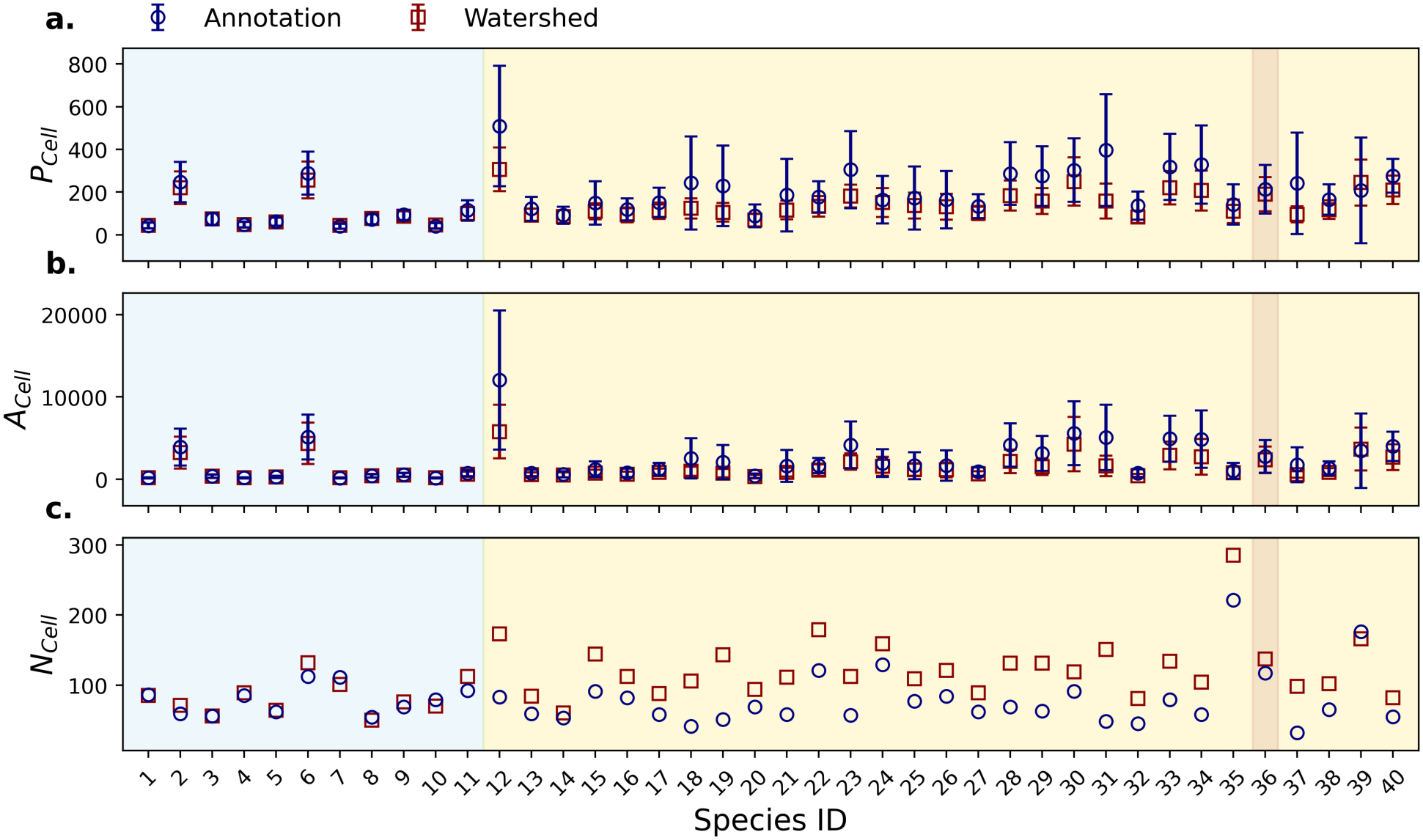
Comparison of watershed and manual segmentation statistics. Comparison of pixel-based statistics derived from watershed and manual cell segmentation of leaf paradermal slices across 40 plant species, ordered by increasing mean arm length, for: **a**. average cell perimeter (*P*_*Cell*_), **b**. average cell area (*A*_*Cell*_), and **c**. cell number (*N*_*Cell*_). Error bars in the panels **a** and **b** represent the standard deviations of within-slice mean values. Blue, yellow, and brown shaded regions indicate non-honeycomb, honeycomb, and neither spongy patterns, respectively.

**Figure S8:**
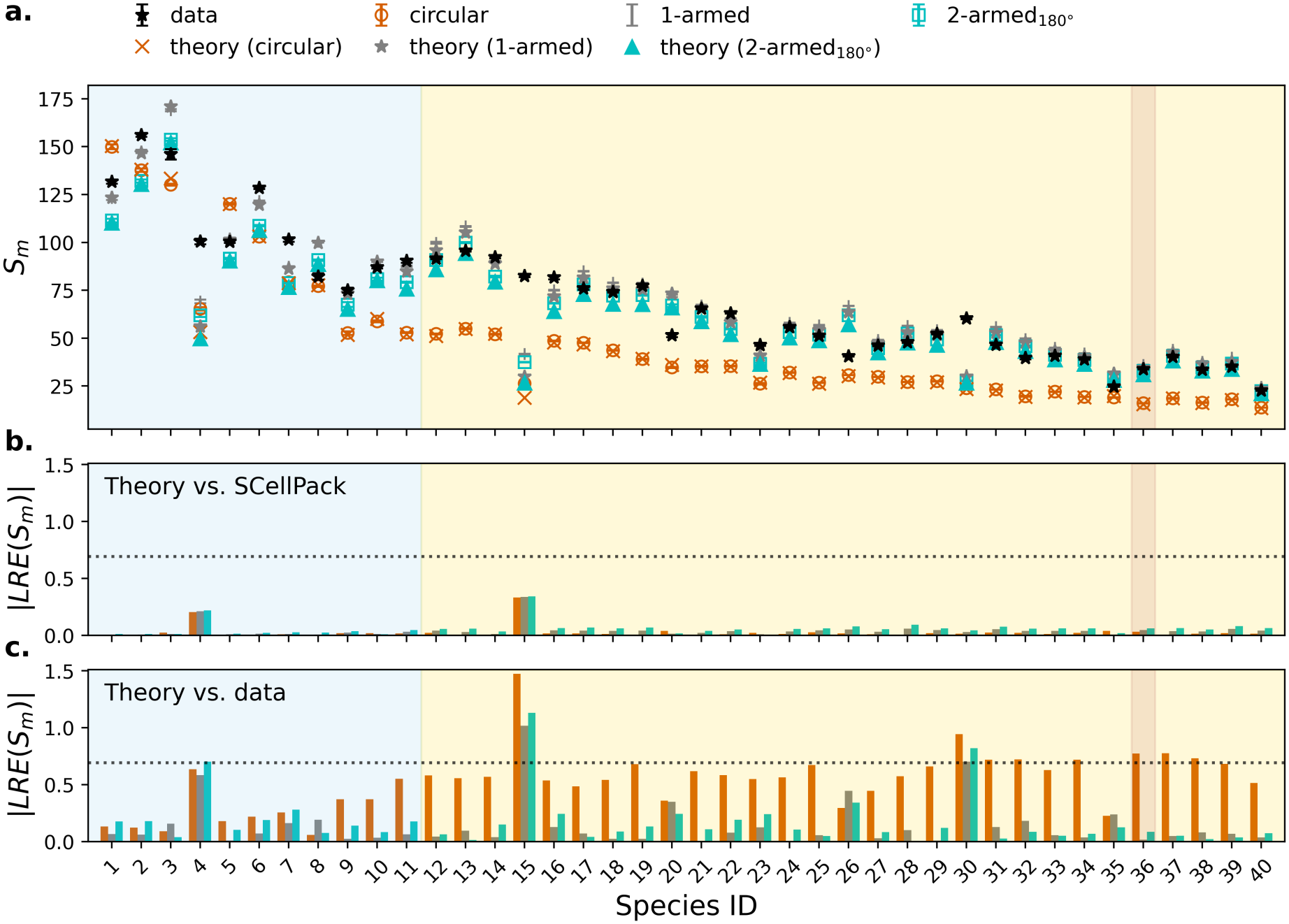
Performance of the Boolean model in predicting mesophyll cell perimeter exposed to intercellular airspace (*S*_*m*_). a. Comparison of theoretical (Boolean model), measured, and simulated mean *S*_*m*_ across 40 plant species. Each marker corresponds to a different cell geometry (circular, 1-armed, and2-armed 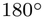). Error bars indicate the standard errors of the slice-level values. Species are ordered by increasing mean arm length. Blue, yellow, and brown shaded regions indicate non-honeycomb, honeycomb, and neither spongy patterns, respectively. **b**. Absolute logarithmic relative error of *S*_*m*_, |*LRE*(*S*_*m*_) | , between theoretical and simulated values across the 40 species and three cell geometries. Bars indicate the reconstruction errors for the corresponding data points of each geometry in panel **a**. The horizontal dotted line marks a two-fold deviation, |*LRE*| = ln(2). **c**. Same as **b**, but showing |*LRE*(*S*_*m*_) | between theoretical and measured values. Markers and colors are consistent across all panels.

**Figure S9:**
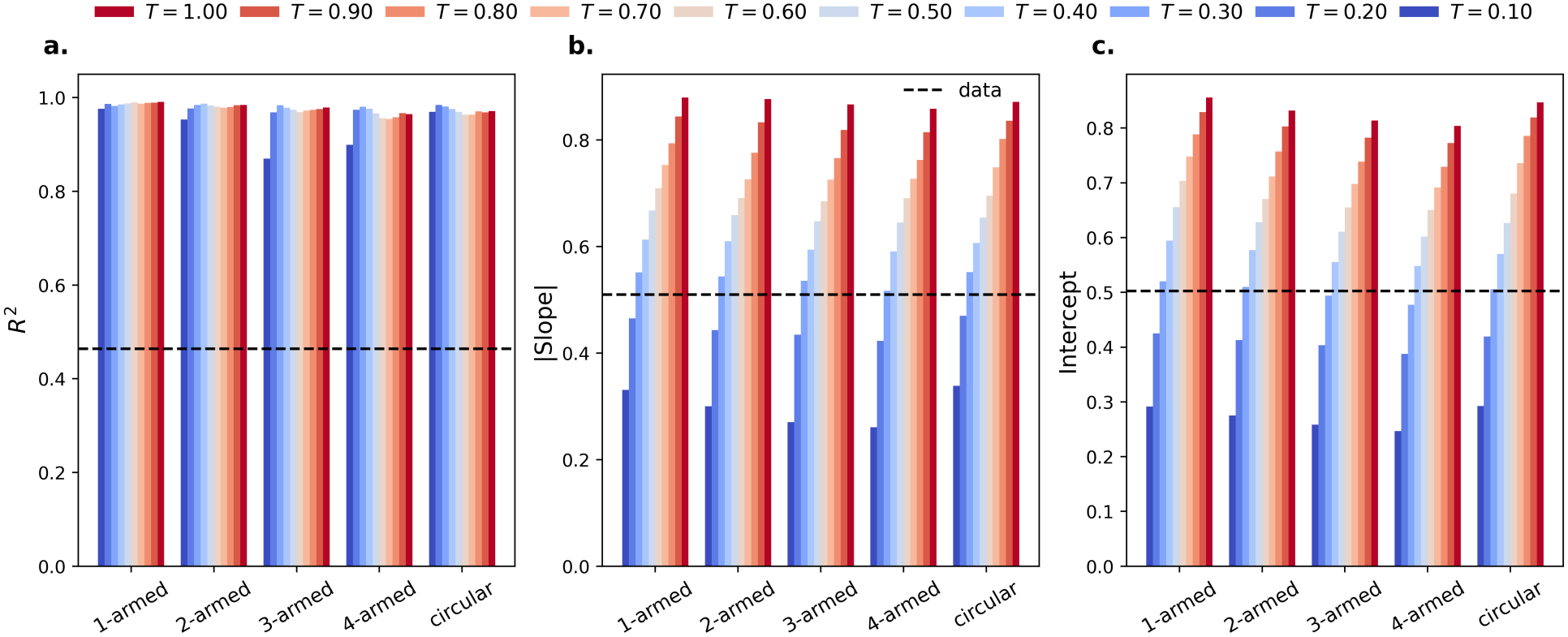
Statistics of the fitted relationship between leaf porosity and shared wall fraction with different truncation fractions. a. Coefficient of determination (*R*^2^), **b**. absolute slope (|*α* |), and **c**. intercept (*m*) of the linear fits between shared wall fraction (*f*_*loss*_) as a function of the fraction of intercellular airspace (*f*_*IAS*_) for different cell geometries across the range of truncation fractions (*T*) tested in SCellPack. Colored bars indicate the fitted values for each *T* , while the dashed black line indicates the corresponding value from the measured data.

**Figure S10:**
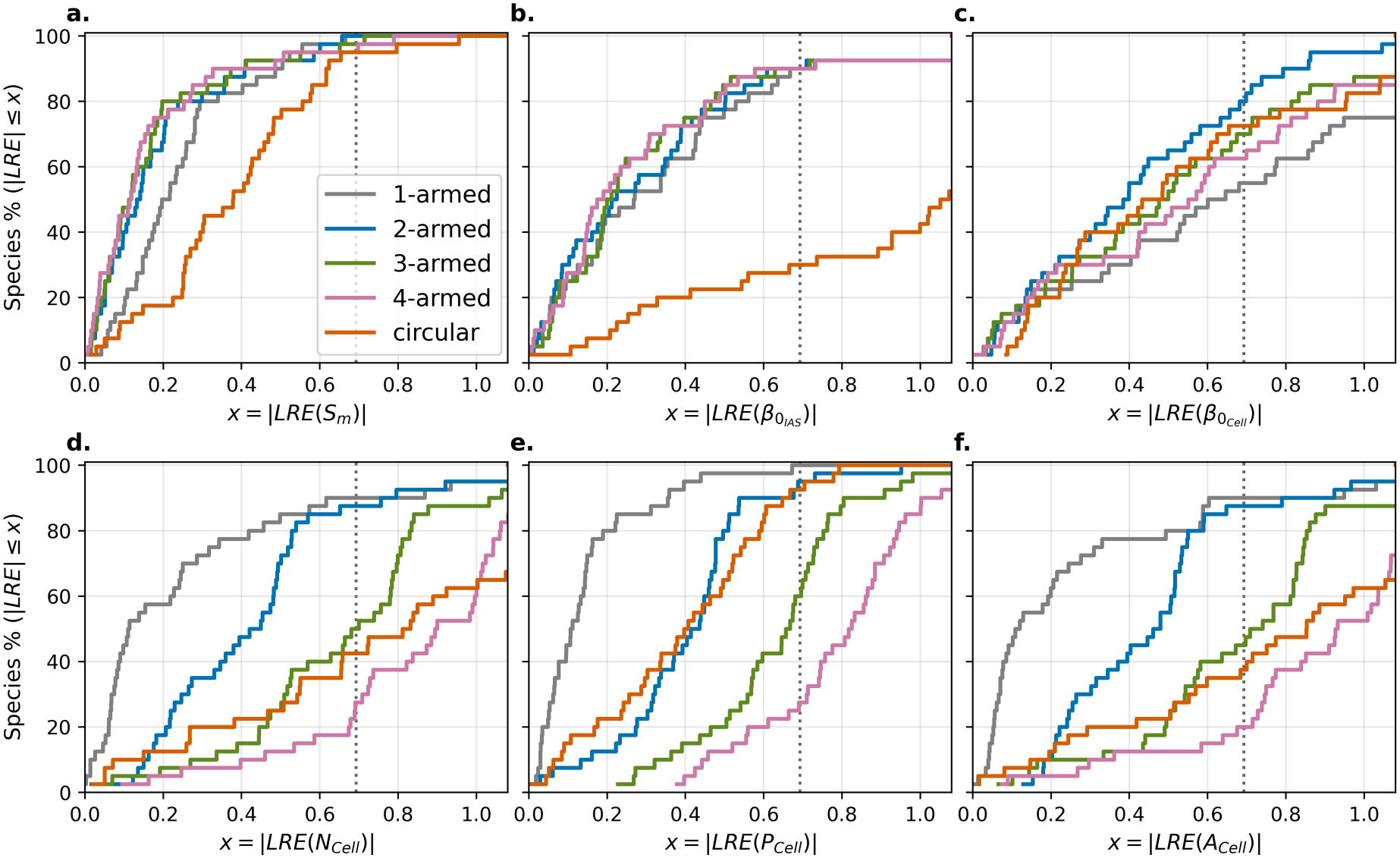
Distributions of SCellPack reconstruction errors for leaf traits with altered truncation fraction (*T*). Cumulative distribution functions of the absolute logarithmic error (| *LRE*|) between simulated and measured values for the traits configured with *T* = 0.4: **a**. cell perimeter exposed to intercellular airspace (*S*_*m*_), **b**. number of intercellular airspace (IAS) connected components 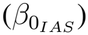, **c**. number of cell connected components 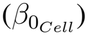, **d**. number of cells (*N*_*Cell*_), **e**. average cell perimeter per leaf area (*P*_*Cell*_), and **f**. average cell area per leaf area (*A*_*Cell*_). Colors indicate different cell geometries (circular, 1-armed, 2-armed, 3-armed, and 4-armed). The curves show the probability that the reconstruction error is less than or equal to a given threshold *x*. The vertical dotted lines mark a two-fold deviation, |*LRE*| = ln(2).

## References

[1] Aleca M Borsuk, Adam B Roddy, Guillaume Théroux-Rancourt, and Craig R Brodersen. Structural organization of the spongy mesophyll. New Phytologist, 234(3):946–960, 2022.

[2] Ichiro Terashima, Yuko T Hanba, Danny Tholen, and Ülo Niinemets. Leaf functionalanatomy in relation to photosynthesis. Plant physiology, 155(1):108–116, 2011.

[3] John R Evans. Mesophyll conductance: walls, membranes and spatial complexity. New Phytologist, 229(4):1864–1876, 2021.

[4] Jaume Flexas, MIQUEL Ribas-Carbo, Antonio Diaz-Espejo, Jeroni Galmés, and Hipólito Medrano. Mesophyll conductance to co2: current knowledge and future prospects. Plant, cell & environment, 31(5):602–621, 2008.

[5] Guillaume Théroux-Rancourt, Adam B Roddy, J Mason Earles, Matthew E Gilbert, Maciej A Zwieniecki, C Kevin Boyce, Danny Tholen, Andrew J McElrone, Kevin A Simonin, and Craig R Brodersen. Maximum co2 diffusion inside leaves is limited by the scaling of cell size and genome size. Proceedings of the Royal Society B: Biological Sciences, 288(1945):20203145, 2021.

[6] Jürgen Knauer, Matthias Cuntz, John R Evans, Ü lo Niinemets, Tiina Tosens, Linda-Liisa Veromann-Jürgenson, Christiane Werner, and Sönke Zaehle. Contrasting anatomical and biochemical controls on mesophyll conductance across plant functional types. New Phytologist, 236(2):357–368, 2022.

[7] Christoph Lehmeier, Radoslaw Pajor, Marjorie R Lundgren, Andrew Mathers, Jen Sloan, Marion Bauch, Alice Mitchell, Chandra Bellasio, Adam Green, Daniel Bouyer, et al. Cell density and airspace patterning in the leaf can be manipulated to increase leaf photosynthetic capacity. The Plant Journal, 92(6):981–994, 2017.

[8] Christopher D Whitewoods. Riddled with holes: Understanding air space formation in plant leaves. PLoS biology, 19(12):e3001475, 2021.

[9] Kimitsune Ishizaki. Development of schizogenous intercellular spaces in plants. Frontiers in Plant Science, 6:497, 2015.

[10] John D Treado, Adam B Roddy, Guillaume Théroux-Rancourt, Liyong Zhang, Chris Ambrose, Craig R Brodersen, Mark D Shattuck, and Corey S O’Hern. Localized growth and remodelling drives spongy mesophyll morphogenesis. Journal of the Royal Society Interface, 19(197), 2022.

[11] Liyong Zhang and Chris Ambrose. Beauty is more than epidermis deep: How cell division and expansion sculpt the leaf spongy mesophyll. Current opinion in plant biology, 79:102542, 2024.

[12] J Mason Earles, Guillaume Théroux-Rancourt, Adam B Roddy, Matthew E Gilbert, Andrew J McElrone, and Craig R Brodersen. Beyond porosity: 3d leaf intercellular airspace traits that impact mesophyll conductance. Plant Physiology, 178(1):148–162, 2018.

[13] Guillaume Théroux-Rancourt, JoséCarlos Herrera, Klara Voggeneder, Federica De Berardinis, Natascha Luijken, Laura Nocker, Tadeja Savi, Susanne Scheffknecht, Moritz Schneck, and Danny Tholen. Analyzing anatomy over three dimensions unpacks the differences in mesophyll diffusive area between sun and shade vitis vinifera leaves. AoB Plants, 15(2):plad001, 2023.

[14] Jeroen DM Schreel, Guillaume Théroux-Rancourt, Pamela K Diggle, Craig Brodersen, and Adam B Roddy. Into the spongy-verse: structural differences between leaf and flower mesophyll. Integrative and comparative biology, 65(1):81–93, 2025.

[15] Joohyun Kang, Kyungyoon Kim, Jooyeon Woo, Thanh Ha Thi Do, and Yuree Lee. Recent insights into air space formation in plant shoots. Current Opinion in Plant Biology, 85:102725, 2025.

[16] Liyong Zhang, Delanie McEvoy, Yen Le, and Chris Ambrose. Live imaging of microtubule organization, cell expansion, and intercellular space formation in arabidopsis leaf spongy mesophyll cells. The Plant Cell, 33(3):623–641, 2021.

[17] Adrienne HK Roeder, Paul T Tarr, Cory Tobin, Xiaolan Zhang, Vijay Chickarmane, Alexandre Cunha, and Elliot M Meyerowitz. Computational morphodynamics of plants: integrating development over space and time. Nature reviews Molecular cell biology, 12(4):265–273, 2011.

[18] Frédéric Boudon, Jérôme Chopard, Olivier Ali, Benjamin Gilles, Olivier Hamant, Arezki Boudaoud, Jan Traas, and Christophe Godin. A computational framework for 3d mechanical modeling of plant morphogenesis with cellular resolution. PLoS computational biology, 11(1):e1003950, 2015.

[19] Bandan Chakrabortty, Ikram Blilou, Ben Scheres, and Bela M Mulder. A computational framework for cortical microtubule dynamics in realistically shaped plant cells. PLoS computational biology, 14(2):e1005959, 2018.

[20] Varsha S Pathare, Nuria Koteyeva, and Asaph B Cousins. Increased adaxial stomatal density is associated with greater mesophyll surface area exposed to intercellular air spaces and mesophyll conductance in diverse c4 grasses. New Phytologist, 225(1):169–182, 2020.

[21] Milad Rahimi-Majd, Alistair Leverett, Arne Neumann, Johannes Kromdijk, and Zoran Nikoloski. Nonlinear models based on leaf architecture traits explain the variability of mesophyll conductance across plant species. Plant, Cell & Environment, 47(12):5158–5171, 2024.

[22] Yusuke Mizokami, Riichi Oguchi, Daisuke Sugiura, Wataru Yamori, Ko Noguchi, and Ichiro Terashima. Cost–benefit analysis of mesophyll conductance: diversities of anatomical, biochemical and environmental determinants. Annals of Botany, 130(3):265–283, 2022.

[23] Sung Nok Chiu, Dietrich Stoyan, Wilfrid S Kendall, and Joseph Mecke. Stochastic geometry and its applications. John Wiley & Sons, 2013.

[24] Stephen B Gray. Local properties of binary images in two dimensions. IEEE Transactions on computers, 100(5):551–561, 1971.

[25] Herbert Edelsbrunner and John Harer. Computational topology: an introduction. American Mathematical Soc., 2010.

[26] Bruno Galerne. Computation of the perimeter of measurable sets via their covariogram. applications to random sets. Image Analysis & Stereology, 30(1):39–51, 2011.

[27] Alice L Baillie and Andrew J Fleming. The developmental relationship between stomata and mesophyll airspace. New Phytologist, 225(3):1120–1126, 2020.

[28] James M Fischer, Guillaume Théroux-Rancourt, Craig R Brodersen, and Andrew DB Leakey. High-resolution microct reveals relationships between stomata and interior leaf anatomy in sorghum. Plant Physiology, page kiag414, 2026.

[29] D Jeulin. Dead leaves models: from space tesselation to random functions. In Proc. of the Symposium on the Advances in the Theory and Applications of Random Sets, pages 137–156. World Scientific, 1996.

[30] Yann Gousseau and François Roueff. The dead leaves model: general results and limits at small scales. arXiv preprint math/0312035, 2003.

[31] Mathew D Penrose. Leaves on the line and in the plane. 2020.

[32] Stefan Van der Walt, Johannes L Schönberger, Juan Nunez-Iglesias, François Boulogne, Joshua D Warner, Neil Yager, Emmanuelle Gouillart, and Tony Yu. scikit-image: image processing in python. PeerJ, 2:e453, 2014.

[33] Pauli Virtanen, Ralf Gommers, Travis E Oliphant, Matt Haberland, Tyler Reddy, David Cournapeau, Evgeni Burovski, Pearu Peterson, Warren Weckesser, Jonathan Bright, et al. Scipy 1.0: fundamental algorithms for scientific computing in python. Nature methods, 17(3):261–272, 2020.

[34] Johannes Schindelin, Ignacio Arganda-Carreras, Erwin Frise, Verena Kaynig, Mark Longair, Tobias Pietzsch, Stephan Preibisch, Curtis Rueden, Stephan Saalfeld, Benjamin Schmid, et al. Fiji: an open-source platform for biological-image analysis. Nature methods, 9(7):676–682, 2012.

